# Exploring the Macroevolutionary Signature of Asymmetric Inheritance at Speciation

**DOI:** 10.1101/2023.02.28.530448

**Authors:** Théo Gaboriau, Joseph A. Tobias, Daniele Silvestro, Nicolas Salamin

## Abstract

Popular comparative phylogenetic models such as Brownian Motion, Ornstein-Ulhenbeck, and their extensions, assume that, at speciation, a trait value is inherited identically by the two descendant species. This assumption contrasts with models of speciation at the micro-evolutionary scale where phenotypic distributions of the descendants are sub-samples of the ancestral distribution. Various described mechanisms of speciation can lead to a displacement of the ancestral phenotypic mean among descendants and an asymmetric inheritance of the ancestral phenotypic variance. In contrast, even macro-evolutionary models that account for intraspecific variance assume symmetrically conserved inheritance of the ancestral phenotypic distribution at speciation. Here we develop an Asymmetric Brownian Motion model (ABM) that relaxes the hypothesis of symmetric and conserved inheritance of the ancestral distribution at the time of speciation. The ABM jointly models the evolution of both intra- and inter-specific phenotypic variation. It also allows the mode of phenotypic inheritance at speciation to be inferred, ranging from a symmetric and conserved inheritance, where descendants inherit the ancestral distribution, to an asymmetric and displaced inheritance, where descendants inherit divergent phenotypic means and variances. To demonstrate this model, we analyze the evolution of beak morphology in Darwin finches, finding evidence of character displacement at speciation. The ABM model helps to bridge micro- and macro-evolutionary models of trait evolution by providing a more robust framework for testing the effects of ecological speciation, character displacement, and niche partitioning on trait evolution at the macro-evolutionary scale.

---

Models describing the evolution of phenotypic traits along phylogenetic trees (also called Phylogenetic Comparative Methods - PCM) are crucial to understanding processes that shaped present biodiversity. Most of these models can be described as multivariate stochastic processes using the phylogenetic tree to describe covariance between species (Lande, 1980b; Felsenstein, 1985). The plethora of new PCM developed in recent years illustrate the importance of these methods in modern macroevolutionary research and allows modelling trait evolution as neutral (Brownian motion, Felsenstein (1973)), drifting towards one or several optima (Ornstein-Ulhenbeck, Hansen (1997); Khabbazian et al. (2016)), varying with a trend (Silvestro et al., 2019) or drifting away from interacting clades (Drury et al., 2016). Models are also able to incorporate evolutionary rates variation through time (Harmon et al., 2010a), among clades (Beaulieu et al., 2012; Castiglione et al., 2018), or with respect to environmental changes (Clavel and Morlon, 2017), other traits (Hansen et al., 2021) or substitution rates (Lartillot and Poujol, 2011).

One underlying and often neglected assumption of these models is the symmetric and complete inheritance of the ancestors’ phenotype by its descendants at a branching event. This assumption is rooted in two characteristics of the PCM. First, phylogenetic trees represent speciation as an instantaneous event in time (Mendes et al., 2018). Second, PCM generally models the evolution of mean phenotypes, ignoring the intraspecific variation. Models incorporating intraspecific trait variance either treat it as a measurement error (Harmon and Losos, 2005) or model its evolution independently of the trait mean (Kostikova et al., 2016; Gaboriau et al., 2020).

Given these structural constraints, it is challenging to consider within the current PCM the progressive nature of the speciation process and its effect on the trait distribution of the descendants. This assumption contrasts with the classic description of speciation at the micro-evolutionary scale (Simpson, 1953; Mayr, 1963; Lande, 1980a; Gavrilets, 2014), in which accelerated trait divergence is often associated with speciation (cladogenetic change). In the context of neutral divergence, descendant species are sub-samples of the parent species population. Differences between incipient species phenotypic distributions are therefore expected by chance (Duchen et al., 2021), especially in the case of peripatric and parapatric speciation where one of the incipient species originates as a small fraction of the parent population size with therefore more chances to diverge from the ancestral trait distribution (Schwämmle et al., 2006; Kopp, 2010). Alternatively, the speciation process can directly cause trait divergence. For instance, dispersing to a new region or environment can increase the probability of speciation because of reduced gene flow and trait divergence by local adaptation and ecological opportunity (Simpson, 1953; Mayr, 1963; Eastman et al., 2013). Island radiations provide striking empirical evidence of this scenario (Losos et al., 2003; Grant and Grant, 2008) with accelerated phenotypic divergence at the time of speciation (Eastman et al., 2013). Other ecological opportunities, such as the extinction of potential competitors or the emergence of a key innovation, are also expected to cause rapid divergence by disruptive selection, leading to sympatric speciation (Gavrilets, 2003; Ackermann and Doebeli, 2004). Given this range of speciation scenarios (Fig. 1), the assumption of equal inheritance of a trait at speciation is likely to be violated, and simulations show this can lead to biased macroevolutionary estimates (Duchen et al., 2021).

**Fig. 1.**
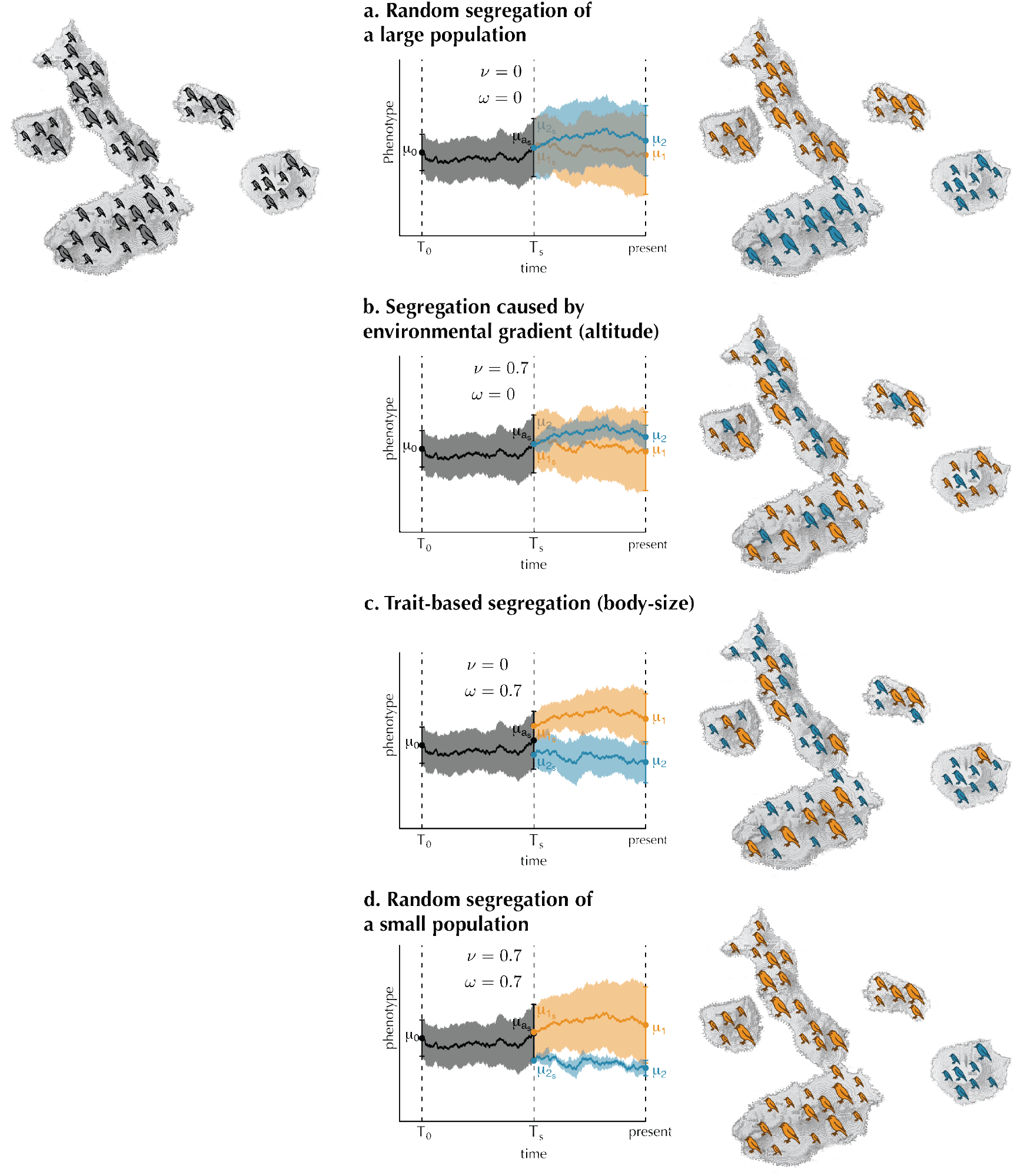
Different scenarios of ancestral distribution’s inheritance is expected depending on the processes that caused segregation. Each line represents a different speciation mechanism from the same ancestral population (top left) with intraspecific variation. The central column represent how ancestral and descendant species distributions change through time. Lines represent the evolution of the mean, and shaded polygons represent the 95% interval of each species distribution. The ancestral is represented in black and the descendants in orange and blue.

Several works proposed testing the Simpsonian evolution tempo at the macroevolutionary scale by modelling anagenetic rates variation (Blomberg et al., 2003; Harmon et al., 2010b) or evolutionary jumps (Landis et al., 2013; Eastman et al., 2013; Duchen et al., 2017). Although they can detect changing rates of phenotypic evolution or phenotypic jumps, those models do not consider cladogenetic changes and might fail to capture its signal. Other attempts proposed modelling the decoupling between anagenetic change and cladogenetic change. The *κ* statistic tests the relationship between branch lengths and evolutionary rates, with *κ* < 1 indicating a faster phenotypic change at speciation than near the tips (Pagel, 1999). Another method proposed to independently model anagenetic and cladogenetic evolutionary rates while incorporating the probability of speciation events masked by extinction, (Bokma, 2008). However, whether these methods identify the signal of cladogenetic changes or capture periods of accelerated anagenetic evolution is still being determined.

An alternative approach to modelling trait evolution uses measurements across multiple individuals per species as input data to jointly infer the evolution of the trait mean across species and of its intraspecific variance (Kostikova et al., 2016; Gaboriau et al., 2020). While this approach unlocks the possibility to model explicitly how the ancestral trait distribution divides between descendent species, its current implementation makes the simplifying assumptions that 1) trait means and variance evolve independently and 2) they identically inherit the ancestral distribution at speciation.

Here, we posit that cladogenetic changes due to asymmetric inheritance at speciation leave a signature in present species trait distribution. Thus, a joint analysis of the evolution of trait mean and variance can reveal the mode of cladogenetic trait inheritance (Fig. 1). We develop a new method to model both cladogenetic and anagenetic changes of phenotypic distribution.

## Materials and Methods

### Model definition

We model the evolution of a phenotypic trait’s intraspecific distribution across a phylogenetic tree with alternative inheritance scenarios at speciation. Specifically, we want to model the situation in which the two incipient species do not inherit the entire phenotypic distribution of the ancestral species. Instead, the process should allow for a continuum between the alternative scenarios presented in Fig. 1. We assume that the trait’s intraspecific distribution is normally distributed. Along branches, we independently model the evolution of the intraspecific phenotypic means and log variances following standard PCM approaches. At the time of a speciation event (*T*_*s*_), represented as a bifurcation in a phylogenetic tree, we first make the hypothesis that the ancestral distribution (*X*_*a*_) shares its 5^*th*^ and 95^*th*^ percentiles respectively with at least one of the descendants’ distributions (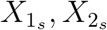, Fig. 1).

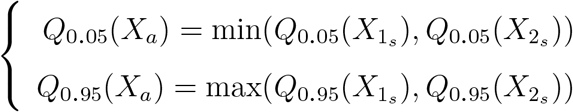

We calculate the 5% and 95% quantiles of a normal distribution (*X* ∼ 𝒩(*μ, σ*)) using the inverse error function (erf^−1^(·)) :

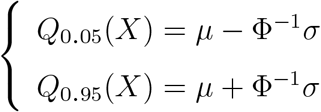

with 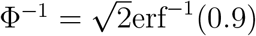

We can define the quantiles of an ancestral species’ distribution as a function of the two descendant species’ variances:

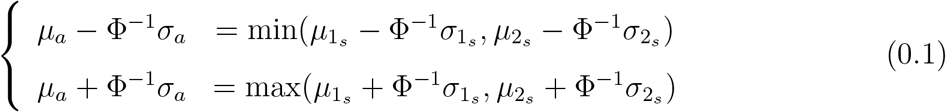

where *μ*_a_ is the mean of the ancestral distribution at the time of speciation, *σ*_*a*_ is the standard deviation of the ancestral distribution at the time of speciation, and 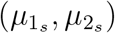 and 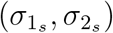 are the means and standard deviations of the descendants’ distributions at the time of speciation. This equation ensures that at least one of the descendants’ distributions respectively shares the 5th and 95th quantiles of the ancestral distribution.

Second, we postulate that the descendants’ distributions might inherit the ancestral variance asymmetrically. Asymmetry between trait variances of the two descendants is denoted by *ν* ∈ [0, 1) as follows:

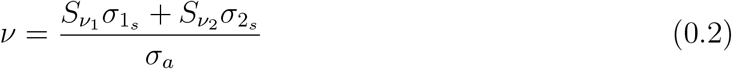

where the switch parameters 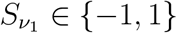 and 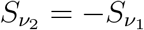 are indicators to specify which descendant inherits the larger part of the ancestral variance. Under this definition, a value of *ν* = 0 indicates a symmetric inheritance of the ancestral variance, while asymmetry grows when *ν* becomes closer to 1. This equation further constrains the difference between the descendants’ variances to be lower than the ancestral variance. This constraint ensures that each descendant’s variance is lower or equal to the ancestral variance.

Third, character displacement might occur at speciation. We measure this process by introducing a parameter *ω* ∈ [0, 1], which represents the displacement between descendants’ mean trait values such that:

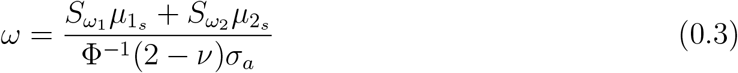

The switch parameters 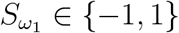 and 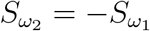 indicate, again, which of the two descendants inherits the highest mean value. Under this definition, *ω* = 0 indicates no displacement between descendants, while *ω* = 1 represents the maximum possible displacement, given Eq. 0.1. The term Ф^−1^(2 − *ν*)*σ*_a_ ensures that the expression of *ω* is compatible with our first hypothesis. If there is asymmetry (*ν* > 0), displacement has to be lower than the ancestral distribution’s 95% interval for the descendants’ distributions 5% and 95% quantiles to remain in that interval. This way, descendants’ means are constrained by the ancestral variance and cannot lie outside the ancestral distribution. These equations (Eqs 0.1, 0.2, 0.3) cover a continuum between alternative inheritance scenarios of the ancestral distribution at speciation (Fig. A2).

### Likelihood of the interspecific distribution

We derive the likelihood of phenotypic means and variances across species given a phylogenetic tree and one or more trait observations per species. We start by showing the calculations on a simple tree with two species sharing a common ancestor. Using Eqn. 0.2 and 0.3, we can express the descendants’ mean and variance as a function of ancestral values, *ν*, and *ω* (see the complete derivation in supplementary methods):

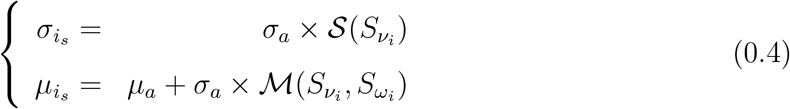

where

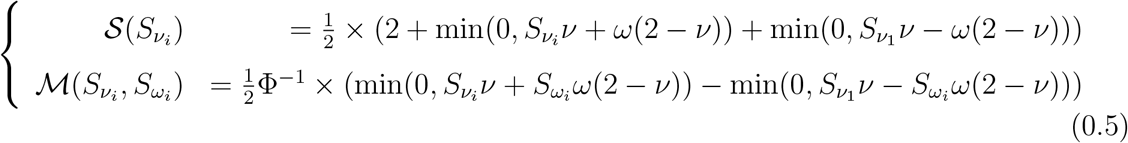

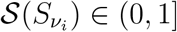 determines the proportion of ancestral variance inherited by descendant i. In the case of asymmetry, 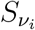 controls whether species i inherits the smaller 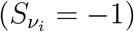 or bigger 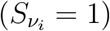 proportion of ancestral variance. Its sister clade will inherit a proportion of ancestral variance equal to 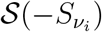. Similarly, 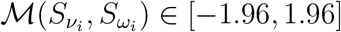 determines the distance between the ancestral mean and descendant *i* mean in terms of ancestral standard deviation units. If 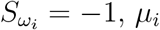 is smaller or equal to the ancestral mean, if 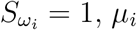 is bigger or equal to the ancestral mean. We consider that means and variances of each species evolve independently along the phylogenetic tree branches following a Brownian motion process (Felsenstein, 1985). We model the evolution of the logarithm of the standard deviation (log(*σ*), hereafter noted *ζ*) and the trait mean following the JIVE algorithm (Kostikova et al., 2016; Gaboriau et al., 2020). At a speciation event, represented by a node in the phylogeny, we inherit descendants’ distributions and obtain the expectations of *ζ*_1_, *ζ*_2_, *μ*_1_ and *μ*_2_ for extant species under this model using Eqn. 0.4:

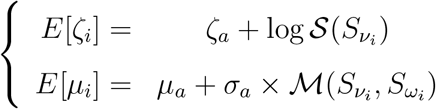

Even though asymmetry and displacement modify expectations for our variables, their variances are not affected by the deterministic process. Thus, we can derive variances from the standard Brownian process:

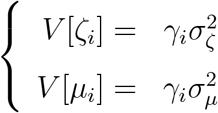

where *γ*_*i*_ is the branch length leading to species *i*, and 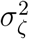 and 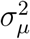 are evolutionary rates of *ζ* and *μ*, respectively. These terms allow us to calculate the probability of *σ*_1_, *σ*_2_, *μ*_1_ and *μ*_2_ given 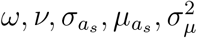 and 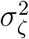 along a phylogenetic tree (Hansen, 1997). For a binary phylogenetic tree with *n* extant species and *n -* 1 ancestral nodes all characterized by their trait distribution *X*_*i*_ ∼ 𝒩(*μ*_*i*_, *σ*_*i*_), we make the simplifying assumption that *ν* and *ω* are constant across every node with a different value for 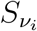 and 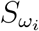. By applying (0.4) to each node and a BM process to each branch, we obtain (see the proof in supplementary methods):

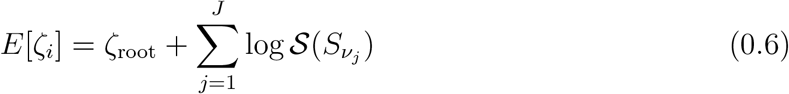

b

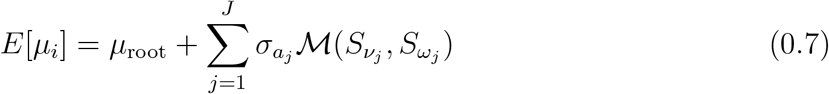

with *J* being the number of branches between the root and species *i* and 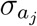 being the standard deviation of *j*’s direct ancestor at speciation. We note that, with fixed *σ*_*i*_, 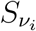 and 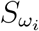, the variance of ***μ*** and ***ζ*** remains constant across nodes, allowing us to use the standard phylogenetic variance-covariance matrix in our calculations. Using *E*[*μ*_*i*_], *E*[*ζ*_*i*_],*V*[*μ*_*i*_] and *V*[*ζ*_*i*_] we can calculate the likelihood functions of ***μ*** and ***ζ*** as multivariate normal distributions (Hansen, 1997).

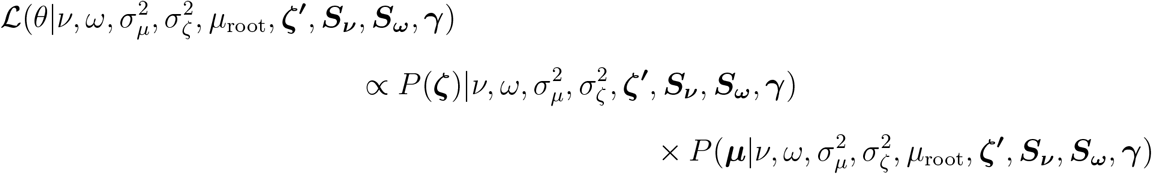

where ***μ*** and ***ζ*** are observed means and log standard deviations. ***ζ*′** and ***γ*** indicate the vector of all ancestral log standard deviations and branch lengths, respectively.

### Parameter estimation

The Bayesian estimation of asymmetry (*ν*), displacement (*ω*) and evolutionary rates 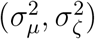 depend on the approximation of ancestral states (***ζ*′**, *μ*_root_) and switch parameters (***S***_*ν*_, ***S***_*ω*_). These parameters can be estimated using Gibbs sampling by calculating their conditional distributions on *ν, ω*, 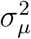 and 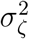. Specifically, we can show that, for any node *k*, the ancestral intraspecific log standard deviation *ζ*_*k*_ is a linear function of its descendants’ *ζ* (see supplementary methods for the proof). Therefore, for every node *k* with *I*_*k*_ extant descendants and *J*_*k*_ descending edges, we have:

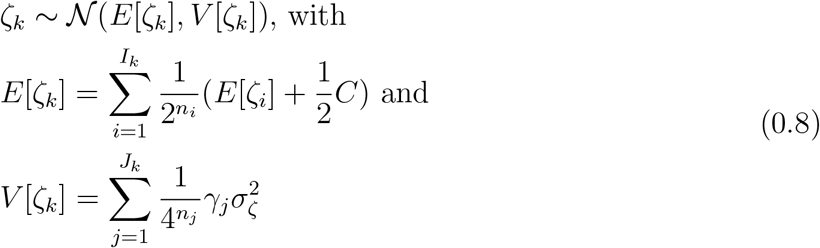

Where *n*_*i*_ and *n*_*j*_ are, respectively, the number of nodes between node *k* and *i, j*, and *C* is a constant for fixed *ω* and *ν*. For every node with direct descendants a and b we can also write:

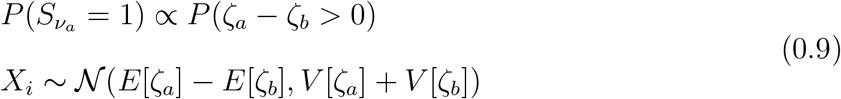

The variable *X*_*i*_ can be used to calculate the conditional probability of 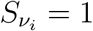.

After the sampling of ***ζ*′** and ***S***_*ν*_ from their respective conditional distributions, we can calculate the conditional distribution of ***μ*′** and ***S***_***ω***_. Similarly, for mean values, we can show that for any node *k, μ*_*k*_ is a linear combination of its descendants’ *μ* (see supplementary methods for the proof). Therefore, for any node *k* with *I*_*k*_ descendants and *J*_*k*_ descending branches, we have:

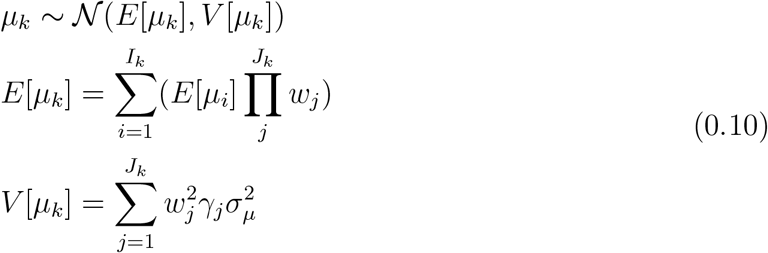

where *w*_*j*_ is a constant for fixed *ω, ν* and 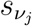 (see proof in supplementary methods). For every node with direct descendants *a* and *b*, we can also write:

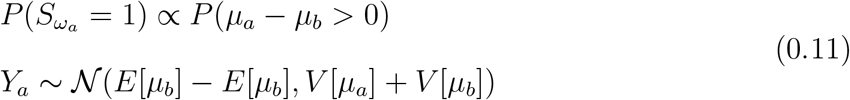

The variable *Y*_*i*_ can be used to calculate the conditional probability of 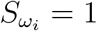. These expressions of conditional distribution for ancestral means, variances, and switch parameters on *ω, ν, μ*_root_, *ζ*_root_), 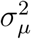 and 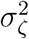, allow us to explore our model’s parameter space using Gibbs sampling. We used multiplier proposals for evolutionary rates and uniform sliding windows *ω, ν* and variance and mean at the root.

### Model validation

We used simulations to test the ability of our model to differentiate between evolutionary processes and to determine whether the estimation of our model’s parameters was accurate. We simulated the random evolution of a trait’s mean and log standard deviation along a set of random phylogenetic trees and simulated asymmetric inheritance at every node with random switch parameters (***S***_*ω*_, ***S***_*ν*_). We simulated five alternative scenarios that represented different evolutionary processes at speciation: (1) Symmetric and Conserved Inheritance (*ν* = 0, *ω* = 0, top left panel of Fig. 1); (2) Symmetric and Displaced Inheritance (*ν* = 0, *ω* = 0.5, top centre panel of Fig. 1); (3) Asymmetric and Conserved Inheritance (*ν* = 0.5, *ω* = 0, bottom left panel of Fig. 1); (4) Asymmetric and Displaced Inheritance (*ν* = 0.5, *ω* = 0.5, bottom right panel of Fig. 1); (5) Intermediate Inheritance (*ν* = 0.2, *ω* = 0.2), bottom centre panel of Fig. 1).

We simulated each scenario on a set of phylogenetic trees containing a different number of species (*n* = *{*20, 50, 100*}*) and different evolutionary rates 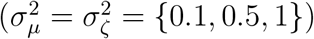. We generated trees using the phytools package based on a birth-death process (Revell, 2012). We simulated phenotypic distributions using the bite package (Gaboriau et al., 2020) with the addition of ancestral distribution’s inheritance process as described above. In total, each set of (*ω, ν, n*, 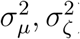) was simulated 100 times leading to 4,500 simulations.

We analyzed simulated datasets by running 500,000 MCMC iterations, sampling every 100 iterations and using uniform priors (𝒰(0.0.99)) for *ω* and *ν* and gamma priors (Γ(1.1, 0.1)) for evolutionary rates. We verified the convergence of the chains using Tracer v1.7.1 (Rambaut et al., 2018) and estimated the means and 95% credible interval of our parameters after removing the first 50,000 iterations as a burn-in. To test whether *ω* and *ν* significantly exceed 0 (under the null hypothesis of a fully symmetric and conserved inheritance), we implemented a Bayesian variable selection algorithm where *ν* and *ω* are multiplied indicators (*I*_*ν*_, *I*_*ν*_ ∈ *{*0, 1*}*) (see Silvestro et al. (2019) and Pimiento et al. (2020) for a similar implementation). The role of the indicators is thus to remove the effect of *ω* or *ν* when set to 0 (in which case the model reduces to a standard Brownian motion), leaving them unaltered when they are set to 1. The value of the indicators is assumed to be unknown and sampled along with the other parameters via MCMC. We set the prior probability to *P*(*I* = 1) = 0.05, meaning that we place a 0.95 probability on a regime with symmetric and conserved inheritance. Based on the posterior sampling frequencies of the indicators, we can estimate Bayes factors to assess the support for models with *ω* > 0 or *ν* > 0 using posterior odds divided by prior odds (Kass and Raftery, 1995).

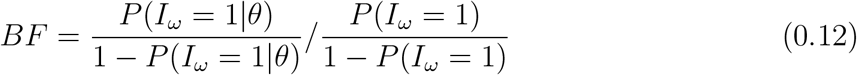

A 2 *log*(*BF*) > 6 supports the relevance of *ω* and *ν* being different from 0 against a symmetric and conserved inheritance (*ω* = *ν* = 0) at speciation.

We also performed a series of control simulations to assess the presence of potential biases. We simulated the effect of an Ornstein-Uhlenbeck model (Hansen, 1997) on *ζ* to test the ability of our model to reject asymmetric and displaced inheritance in a dataset generated by an Ornstein-Uhlenbeck process. We also tested the effect of extant species’ undersampling by resampling the simulated datasets with heterogeneous sample sizes. We resampled five individuals for 25% of the species and 100 individuals for other species and recalculated species means and variances from those samples. Finally, we simulated the effect of process heterogeneity along the tree. We mapped regimes along the trees using two alternative methods. We generated the first dataset by simulating the evolution of a binary trait with symmetrical transition rates. We generated three datasets with different transition rates *q* = *{*0.2, 0.5, 0.8*}*. The binary trait represented a transition between classic BM and ABM regimes. The second dataset was generated by randomly assigning regimes to the tree’s nodes. We generated three datasets with different proportions of nodes under a classic BM regime *p* = *{*0.2, 0.5, 0.8*}*. For all these control simulations, we ran MCMC chains and estimated Bayes Factors and parameters as described above.

### Application

Ecologists and evolutionary biologists often use Darwin’s finches and other Coerebinae to illustrate adaptive radiation driven by character displacement (Grant and Grant, 2008). From a single granivore ancestor, the Coerebinae rapidly diversified in the Galapagos into a diverse clade and covered as many diets as the rest of the Thraupidae family (Reaney et al., 2020) thanks to their considerable diversity of beak shapes. Previous works often link the ecological opportunity brought by the colonization of oceanic islands with this rapid diversification (Grant and Grant, 2008; Reaney et al., 2020; Tobias et al., 2020). However, it remains unclear whether the speciation process is associated with accelerated evolutionary rates of beak shape caused by competition or whether it remained constant during Coerebinae diversification (Tobias et al., 2020). We use the ABM model to test for accelerated evolutionary rates at speciation in Coerebinae. We obtained data for the beak morphology of Coerebinae’s individuals from each extant species (total culmen, beak nares, beak depth and beak width, Fig. 2) (Pigot et al., 2020) and a time-calibrated phylogeny of the clade (*n* = 14)(Burns et al., 2014). We ran two independent MCMC chains for 2,000,000 iterations, sampling every 1,000 iterations for each trait. To determine whether our data shows a signal of asymmetric or displaced inheritance, we employed the variable selection approach for *ν* and *ω* and estimated Bayes factors. We ran two independent chains with fixed indicators for parameter estimation according to the model selection procedure described above. We then calculated mean posterior estimates of our parameters after removing an appropriate burn-in.

**Fig. 2.**
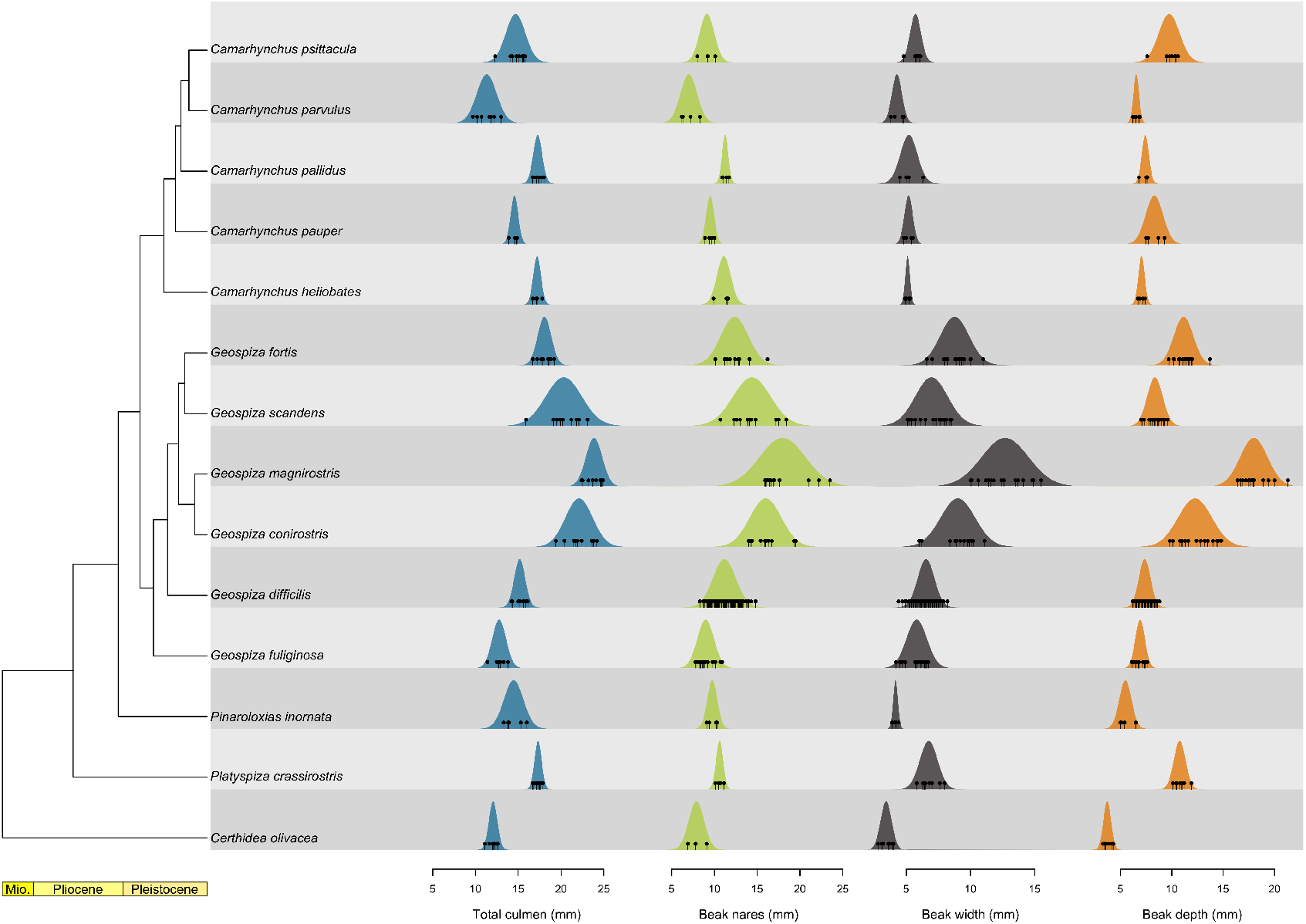
Beak morphology distributions per species in Coerebinae. The left represents the time-calibrated phylogeny of Coerebinae and the right panels represent estimated trait distributions from individual observation (dots) for each species and each trait used in the analysis

## Results

### Performance of the ABM model

Our simulations showed that the signal of different modes of cladogenetic trait inheritance, including trait asymmetry and displacement, can be correctly identified with the ABM model using Bayes factors tests. While most analyses reached convergence, about 20% struggled to yield high ESS values. Indeed we found that for simulations with high evolutionary rates 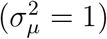, 40% of the chains failed to reach convergence with many fluctuations on evolutionary rates’ estimation (Fig. A14). These non-convergent runs often lead to overestimating evolutionary rates and rejecting asymmetric or displaced inheritance.

Our simulations showed type I and type II error (hereafter indicated with *α* and *β*, respectively) lower than 0.05 at low evolutionary rates for both *ν* and *ω* (Fig. 3).

**Fig. 3.**
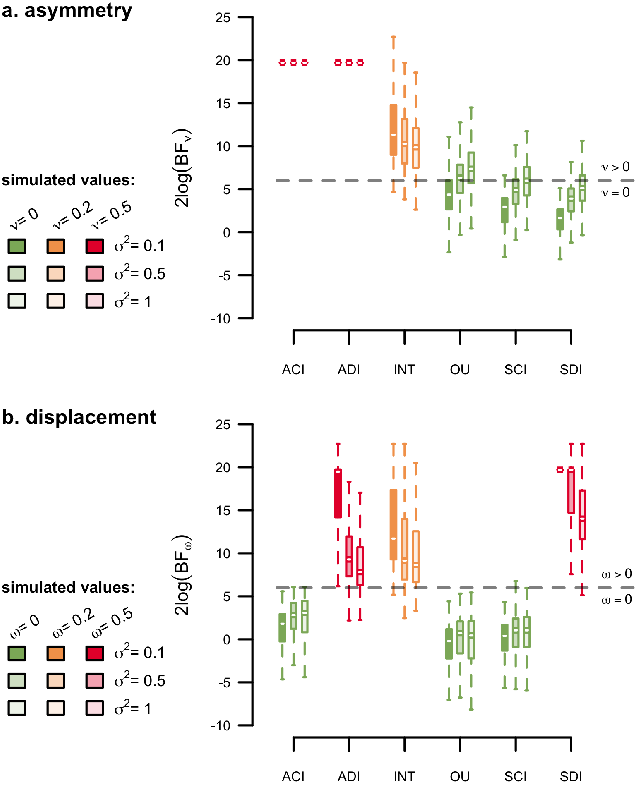
Variable selection results on the simulated datasets. The upper panel represents the 2*log*(*BF*) of displaced vs conserved inheritance in function of the simulated scenarios. Most simulations involving displacement led to a *log*(*BF*_*ω*_) > 6. The lower panel represents the 2*log*(*BF*) of asymmetric vs symmetric inheritance in function of the simulated scenarios. Most simulations involving asymmetry led to a log(*BF*_*ω*_) > 6. Simulations with high evolutionary rates led to less accurate model selection. ACI : Asymmetric and Conserved Inheritance (*ν* = 0.5, *ω* = 0); ADI : Asymmetric and Displaced inheritance (*ν* = 0.5, *ω* = 0.5); INT : Intermediate scenario (*ν* = 0.2, *ω* = 0.2); OU : SCI with Ornstein-Ulhenbeck on the *ζ* (*ν* = 0, *ω* = 0); SCI : Symmetric and Conserved Inheritance (*ν* = 0, *ω* = 0); SDI : Symmetric and Displaced Inheritance (*ν* = 0, *ω* = 0.5).

Simulations with symmetric inheritance (*i*.*e. ν* = 0) showed a slightly higher type II error (*β* < 0.1) for *ν* even with low levels of asymmetry and high evolutionary rates. However, the model was less reliable in consistently rejecting asymmetry in simulations with *ν* = 0 and high evolutionary rates (*α* = 0.31). Error rate further increased in the case of stabilizing selection for intraspecific variance (OU model, *α* = 0.62), an evolutionary mode currently not implemented in the ABM framework. Simulations in the absence of displacement (*i*.*e. ω* = 0) presented a low type I error for *ω*, while results for *ω* > 0 were more contrasted. In the case of symmetric inheritance (*ν* = 0), displacement was correctly identified, while higher levels of asymmetry increased the chances of incorrect rejection of displaced inheritance for high evolutionary rates (*β* = 0.14). The number of tips had a negligible effect on model identification. In conclusion, model testing through Bayes factors correctly identified instances of cladogenetic asymmetries and displacement (or their absence) in cases of relatively low evolutionary rate. In contrast, high evolutionary rates appeared to weaken the signal.

The accuracy of parameter estimation varied depending on the simulation scenarios, but in most cases, the estimation of *ω, ν* and root state was accurate and unbiased (Fig. 4). The accuracy of parameter estimation decreased with increasing evolutionary rates leading to either over or under-estimation depending on the parameter. In particular, we observed that *ω* tends to be underestimated for high levels of displacement, asymmetry and evolutionary rates. We also observe more correlation between variables in the MCMCs ran with this scenario (Fig. A15, A17, A16, A18). With high evolutionary rates, it becomes impossible to differentiate whether a fast change in phenotypic distribution is due to anagenetic or cladogenetic changes. Additionally, in the case of stabilizing selection for the intraspecific variance (OU), we observed an overestimation of *ν* and 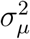. Here again, the number of tips did not affect parameter estimation. The Gibbs sampling procedure accurately estimates ancestral states and switch parameters at the tree’s internal nodes, despite increased uncertainties for simulations with high evolutionary rates.

**Fig. 4.**
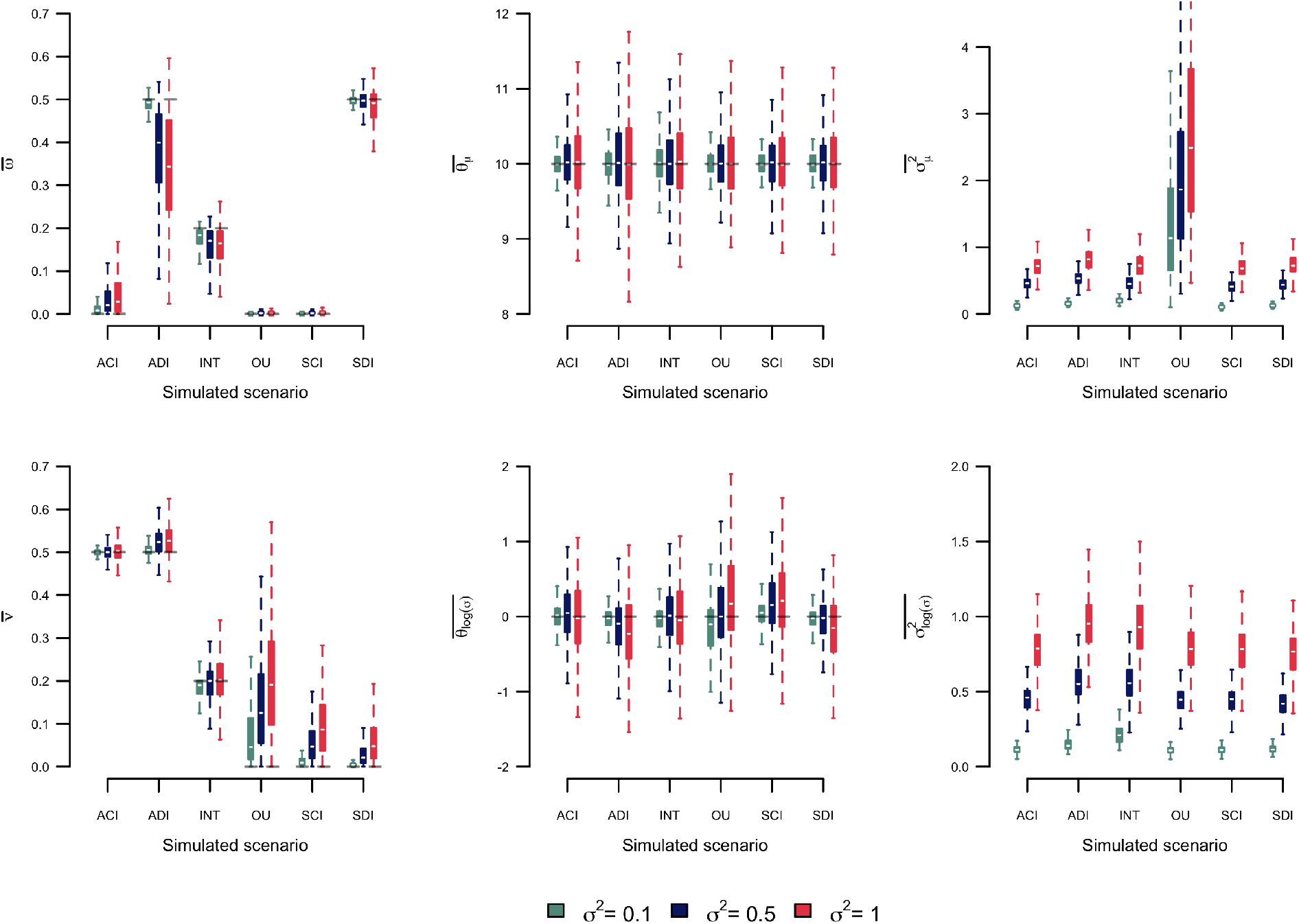
Parameter estimation on the simulated datasets. Each panel represent mean parameter estimation in function of the simulated scenario. The grey dashed lines represent the parameter value used to simulate the dataset. ACI : Asymmetric and Conserved Inheritance (*ν* = 0.5, *ω* = 0); ADI : Asymmetric and Displaced inheritance (*ν* = 0.5, *ω* = 0.5); INT : Intermediate scenario (*ν* = 0.2, *ω* = 0.2); OU : SCI with Ornstein-Ulhenbeck on the *ζ* (*ν* = 0, *ω* = 0); SCI : Symmetric and Conserved Inheritance (*ν* = 0, *ω* = 0); SDI : Symmetric and Displaced Inheritance (*ν* = 0, *ω* = 0.5).

### Coerebinae evolution

The ABM model detected consistent displaced inheritance for three of the four beak trait distributions with log BF comparing models with *ω* = 0 or *ω* > 0 being higher than 6, indicating strong support for cladogenetic displacement (Tab. 1). Specifically, we found evidence for displaced inheritance of total culmen, bill nares and bill depth, although the estimated values of *ω* for the first two were low. In contrast, we found a consistent rejection of asymmetric inheritance for all traits with negative log BF values indicating strong support for *ν* = 0 (Tab. 1). We also observe values of similar magnitude across all traits for the estimates of *μ*_root_, 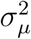 and 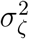, while the estimated *ζ*_root_ are much lower for bill width than for any other traits relative to observed *ζ* in present species.

**Table 1.** Results of the ABM model fitted on Darwin’s finches beak shape traits. The log(BF) columns represent the results of MCMC chains ran with the variable selection algorithm. Estimated parameters (mean posterior) are calculated from MCMC chains ran with selected variables from the previous approach.

| Variable | $2\log(BF_\omega)$ | $2\log(BF_\nu)$ | $\bar{\omega}$ | $\bar{\nu}$ | $\mu_{root}^-$ | $\sigma_{root}^-$ | $\bar{\sigma}_\mu^2$ | $\bar{\sigma}_\zeta^2$ |
| --- | --- | --- | --- | --- | --- | --- | --- | --- |
| Total culmen | 6.24 | -4.35 | 0.09 | - | 14.53 | 13.60 | 0.96 | 0.37 |
| Bill nares | 6.14 | -4.78 | 0.08 | - | 9.33 | 10.75 | 0.67 | 0.31 |
| Bill width | 1.10 | -6.19 | - | - | 5.06 | 0.55 | 1.19 | 0.17 |
| Bill depth | 6.32 | -5.25 | 0.91 | - | 6.36 | 14.93 | 1.18 | 0.42 |

## Discussion

Evolutionary processes are all constrained by the ability of individuals to transmit their genes to the next generation. Most forces affecting evolution (*e*.*g*. selection) therefore unfold their effects at the individual level (Lande, 1976; Hallgrímsson and Hall, 2005; Kaliontzopoulou et al., 2018). Nevertheless, current phylogenetic comparative methods attempting to estimate these forces at a macro-evolutionary scale make the simplifying assumption that species, not individuals, are the fundamental unit of evolutionary mechanisms driving trait evolution. This assumption is rounded on reducing mathematical complexity rather than on theoretical expectations and can lead to biased estimates (Duchen et al., 2021). Using the species as the unit of phenotypic variation does not allow for considering drift, mutation and recombination, which are thought to be the main forces generating variation at the microevolutionary scale (Mayr, 1963; Lande, 1976) and recent methodological developments showed that incorporating intraspecific trait variance in the model can substantially improve our understanding of traits within and among species (Kostikova et al., 2016; Gaboriau et al., 2020). However, even these models assume that speciation is an instantaneous event and that the ancestral lineage’s full range of trait values is passed to the descendants without modifications. This assumption limits the possibility of testing for the effect of divergence on trait evolution (Schluter, 2000; Turelli et al., 2001; Bokma, 2008; Duchen et al., 2021).

In this study, we presented a model that relaxed some of these assumptions, providing a framework to infer how a trait (and its intraspecific variation) may be asymmetrically inherited by descendent species, reflecting different magnitudes of trait displacement. Even though our model is not individual-based, it incorporates multiple measurements per species. It allows us to model the evolution of a trait mean and variance, thus approximating intraspecific variation.

It thus studies the same system as micro-evolutionary models but does it at a different scale. Results from our ABM can then capture the signal of different mechanisms of trait segregation at speciation, such as trait-based segregation, random segregation or segregation caused by environmental gradients. In turn, experts can use their knowledge regarding such mechanisms as prior information for the ABM. Second, the ABM considers the effect of ancestral intraspecific variation on character displacement between descendants at speciation. As such, estimated cladogenetic changes are constrained and represent realistic mechanisms. The ancestral distribution inheritance process allows the modelling of fast character displacement realistically in the light of the modern synthesis instead of considering stochastic cladogenetic jumps. It reproduces the effect of several micro-evolutionary processes associated with the speciation process. For instance, character displacement with low overlap between descendants distribution can be the effect of disruptive selection associated with assortative mating (Dieckman et al., 2004; Seehausen and Van Alphen, 1999; Bolnick, 2001; Gavrilets, 2003; Dijkstra and Border, 2018; Tobias et al., 2014). Alternatively, the isolation of small populations following a colonisation event can lead to an asymmetric inheritance of the ancestral variance. In turn, local adaptation can cause rapid evolution of remote populations leading to significant displacement (Simpson, 1953; Losos and Ricklefs, 2009; Wagner et al., 2012; Mahler et al., 2013). In this scheme, the ancestral variance does not represent the realised variance of ancestral populations but more the evolvability of ancestral species (Wagner and Altenberg, 1996; Abzhanov, 2017; Payne and Wagner, 2019), meant here as the genetic potential to create diversity as a response to selection drift and recombination. Because evolvability depends on genetic and structural constraints (Pigliucci, 2008), it can be seen as a heritable trait and modelled as a diffusion process (Kostikova et al., 2016; Gaboriau et al., 2020). Furthermore, speciation events associated with character displacement and specialisation can reduce the evolvability of descendant species and lead to the realised phenotypic variance that we observe today.

### Model performance

Our simulations suggest that the ABM model can identify homogeneous regimes of cladogenetic asymmetric inheritance of the variance and displaced inheritance of the mean. Our variable selection algorithm allows rejecting the hypothesis of asymmetry (*ν*) or displacement (*ω*) when their signal is absent or weak. However, high anagenetic evolutionary rates can mask the signal of asymmetric or displaced inheritance, leading to model misidentifications. The rejection of displacement is conservative, meaning that the ABM model does not typically find spurious evidence. In contrast, the signal of asymmetric inheritance bears the same signal as high evolutionary rates. However, rates of anagenetic evolution that mask the signal of symmetric and displaced inheritance are high and generate the same signal as white noise, erasing the covariance between species. We also show that the effect of stabilising selection on the log variance, modelled here as an OU process, undersampling and heterogeneous process, also decreases the power of our model while increasing the uncertainty ranges in the estimated parameters.

We also found that the number of tips considered has a low effect on the parameter estimation or variable selection procedure. We base this observation on simulations of a constant *ν* and *ω* across all nodes. With empirical datasets, the chances that this hypothesis is violated increase with the number of tips, as heterogeneous evolutionary regimes are more likely in larger datasets. However, it is essential to note that standard PCMs also make the hypothesis that trait inheritance at speciation is homogeneous across all nodes and that it is symmetric and conserved. As such, the ABM improves the standard BM process and opens promising perspectives in integrating individual-level processes in macro-evolutionary analyses.

### Phenotypic evolution in Darwin’s finches

The ABM model identified consistent character displacement and variance partitioning in traits associated with beak shape in Darwin’s finches. This finding indicates that speciation in this well-known group of birds is associated with cladogenetic divergence of beak shape, likely linked to niche partitioning (Felice et al., 2019). It also demonstrates that the intra- and interspecific variation in Darwin’s finches’ beak shape is not only driven by anagenetic changes but is also the result of local adaptation happening simultaneously with allopatric speciation (Tobias et al., 2020) or divergence driven by competition or other interactions. This pattern is consistent with previous studies finding an association between elevated speciation rates and fast morphological evolution in Darwin’s finches (Reaney et al., 2020). It further provides indirect evidence of the link between speciation and niche partitioning at the macroevolutionary scale for this clade.

Although the estimated values for *ω* might appear small, the displacement of descendants’ mean phenotypes during speciation represents a ≈ 10% of the ancestral 95% interval (or more if we consider the frequent underestimation of *ω* detected in our simulations). It is thus likely that displacement and variance partitioning are higher than estimated. We also observed that estimated variances at the root are much higher for the three traits under a regime of *ω* > 0. This difference comes from the assumption of the ABM model that the regimes are constant through all nodes. A model with *ω* > 0 or *ν* > 0 thus assumes that the intraspecific variance divides at each node. The high morphological variation in Darwin’s finches has often been associated with their high cranial shape modularity (Tokita et al., 2017; Abzhanov, 2017) and the high variability in their development (Mallarino et al., 2012). The high ancestral variance estimated can thus be associated with that evolutionary potential (*i*.*e*. evolvability) progressively constrained by competitive interactions every time a new species arises.

### Future improvements to the ABM model

With the current implementation, the ABM only considers constant and neutral anagenetic evolution. Allowing other existing modes of anagenetic evolution (time-variation, selective optima, density dependence, evolutionary trend) would increase the flexibility of the model and could tackle some issues, such as the effect of stabilising selection on the log variance and the underestimation of *ω*. The ABM model also assumes currently homogeneous regimes of cladogenetic trait inheritance. This assumption is unlikely to hold in nature as many factors are involved in the speciation processes that can generate heterogeneous patterns of trait distribution inheritance at the time of speciation. Therefore it can only identify broad tendencies without being precise about that heterogeneity. One straightforward solution to circumvent this issue could be to allow for several regimes of cladogenetic trait distribution inheritance in the same way as existing comparative methods. Those regimes could be introduced *a priori* using alternative knowledge about the clades evolution (Beaulieu et al., 2012; Zhang et al., 2022) or found based on the data (Uyeda and Harmon, 2014). Alternatively, we could extend this model to allow for the dependency between *ν* and *ω* on predictor variables. For instance, the effect of present geographical overlap could be used as a predictor of character displacement, while species densities could predict asymmetry.

Finally, the model assumes that every speciation event is observed in the phylogeny, ignoring the effect of extinction. Incorporating speciation and extinction rates and the effect of hidden speciation events would likely improve the predictions of the ABM model (Bokma, 2008). Furthermore, the asymmetric inheritance process represents a discrete realisation of a continuous event: speciation. The asymmetric and displaced inheritance described by the model is thus an abstraction of a continuous process of divergence from the ancestral distribution into two independent descendant distributions. This abstraction has some limitations, as it assumes that every speciation event happens at the same pace. This assumption can be problematic as we expect a causal link between descendants’ distributions divergence and time to complete speciation. Introducing the concept of protracted speciation (Rosindell et al., 2010; Hua et al., 2022) in the ABM framework would allow taking into account the gradual nature of the speciation process and model a gradual divergence of daughter species distributions. However, the identification of this process might be difficult.

Overall, the ABM brings comparative methods one step closer to integrating individual variation into macro-evolutionary models. This is timely because phenotypic and phylogenetic datasets are growing in size and completeness, making it increasingly easy to obtain information about individual variation in traits for entire clades (see Tobias (2022); Schleuning et al. (2023)). The unprecedented accessibility of individual-level data across large numbers of species gives us the opportunity to incorporate individual variation at the macro-evolutionary scale. It highlights the need for a new generation of models designed to test individual-level predictions. By addressing one aspect of this challenge, we hope that the ABM can inspire further progress in developing the models required to explore emerging patterns in macroevolutionary diversification.

## Acknowledgments

Data available from the Dryad Digital Repository: https://doi.org/10.5061/dryad.q573n5tns

We declare that we have no conflict of interest to disclose.

## Appendix

### Likelihood of the interspecific distribution

The goal of this section is to estimate the likelihood of the means and variances of observed species given a phylogenetic tree. We start with a simple tree with two species sharing a common ancestor. Using (0.2) and (0.3) we can express the variance and mean of the descendants as a function of each other:

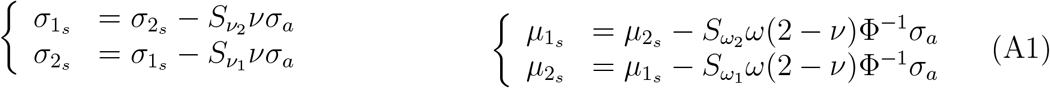

Combining (A1) with (0.1) we get:

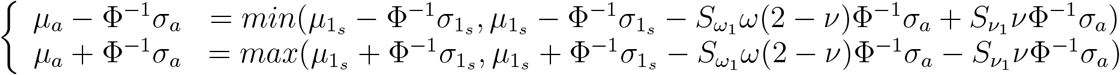

which simplifies to:

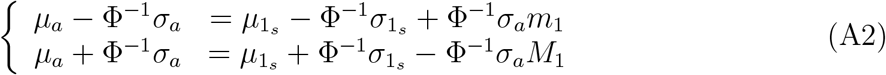

with 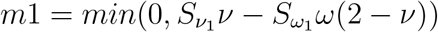 and 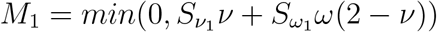. Solving this set of equations for 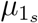 and 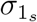 gives us an expression of the first descendant’s mean and variance in a function of *ν, ω*, 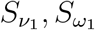:

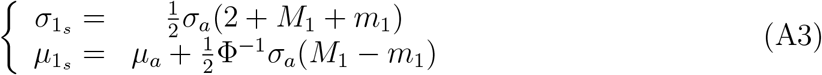

with the same method for species 2 we get:

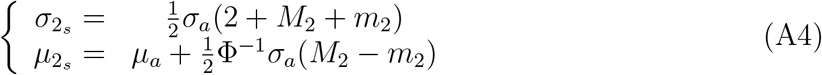

with 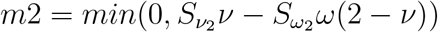 and 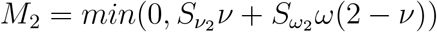.

We note :

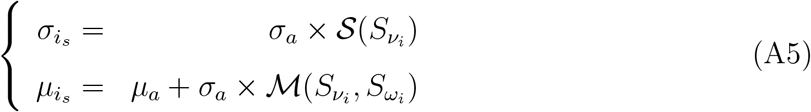

where

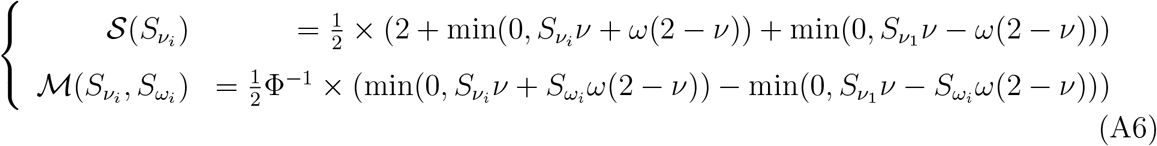

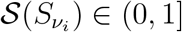 determines the proportion of the ancestral variance inherited by descendant *i*. In the case of asymmetry, 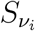 controls whether species i inherits the smaller 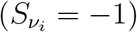 or bigger 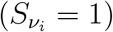 proportion of the ancestral variance. Its sister clade will inherit a proportion of the ancestral variance equal to 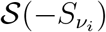. Similarly,

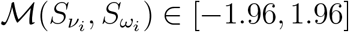 determines the distance between the ancestral mean and descendant *i* mean in terms of ancestral standard deviation units. If 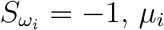 is smaller or equal to the ancestral mean, if 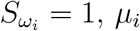 is bigger or equal to the ancestral mean.

We consider that species means and variances evolve independently along branches following a Brownian motion. We model the evolution of the logarithm of the standard deviation (*ζ*) for the variance following the JIVE algorithm; At each node, we apply the asymmetric and displaced inheritance process described above. We can calculate the expectations of *ζ*_1_, *ζ*_2_, *μ*_1_ and *μ*_2_ at present according to this model using (A3 and A4):

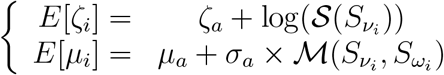

Even thought it modifies the expectations for our variables, the ancestral distribution inheritance process does not affect their variances for fixed *ν* and *ω*. We thus have standard variances derived from the BM process:

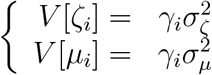

*γ*_*i*_ being the length of the terminal branch leading to species *i* and 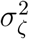 and 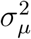 being respectively the evolutionary rates of *ζ* and *μ*. We can then calculate the probability of *σ*_1_, *σ*_2_, *μ*_1_ and *μ*_2_ given *ω, ν*, 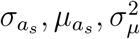 and 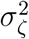 and obtain the likelihood of the tree.

**Fig. A1.**
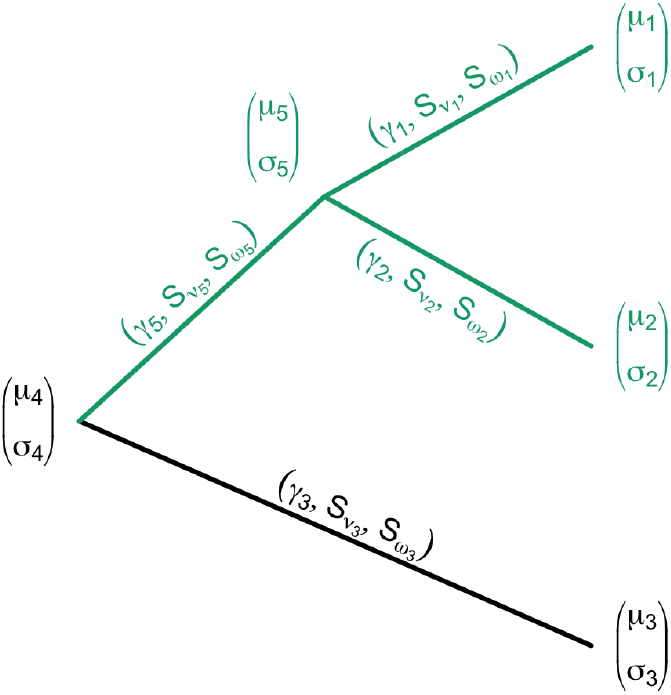
Summary of the notations on the phylogenetic trees: Extant species phenotypic distribution is represented at the tips, ancestral species distribution at the time of speciation is represented at the nodes, branch lengths and inheritance switch parameters are represented along branches. The clade in green represents the initial species complex used to present the ancestral distribution inheritance process.

To calculate the variances and expectations for a slightly more complicated tree (Fig. A1) we make the hypothesis that *ν* and *ω* are constant across every node while we allow every node to have a different value for 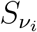 and 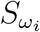:

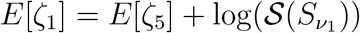

with

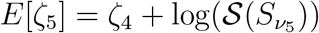

leading to

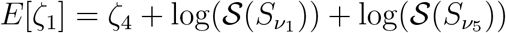

Similarly we have

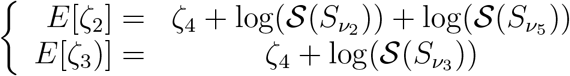

As mentioned earlier, the variances are not affected by the inheritance process and are the standard variances of a BM process:

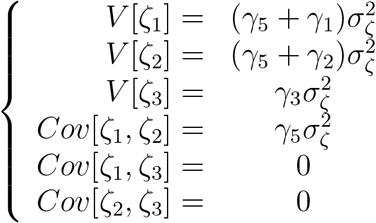

Following the same method, we can find expressions *E*[*μ*_*i*_] as a function of *μ*_4_, *ω, ν* and *S*_*ν*_, *S*_*ω*_ for every node.

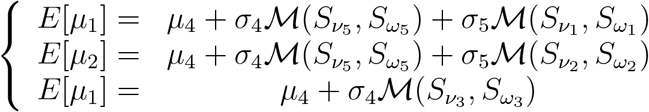

and

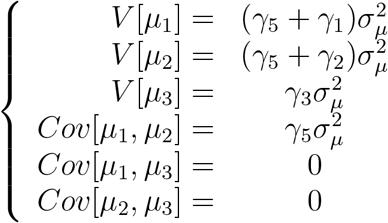

Now if we consider a dichotomous phylogenetic tree with n extant species and n *–* 1 ancestral species all characterized by their trait distribution *X*_*i*_ ∼ 𝒩(*μ*_*i*_, *σ*_*i*_). Every extant and ancestral species, except the root species, is also characterized by switch parameters 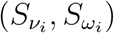 that controls how it inherited its ancestral distribution at the time speciation and by the length of its ascending branch *γ*_*j*_ (Fig. A1).

By applying (A3) and (A4) to each node and a BM process to each branch we get :

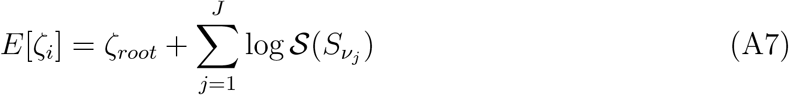

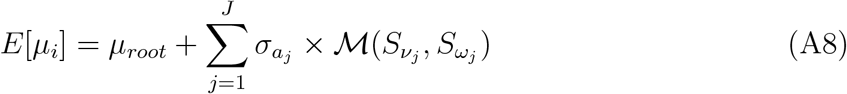

with J being the number of branches between the root and species *i* and 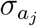 being the standard deviation of the direct ancestor of j at the time of speciation. However, with fixed *σ*_*i*_, 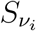 and 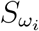, the variance of ***μ*** and ***ζ*** remain constant across nodes, allowing the use of a standard phylogenetic variance-covariance matrix. Using *E*[*μ*_*i*_], *E*[*ζ*_*i*_],*V*[*μ*_*i*_] and *V*[*σ*_*i*_] we can calculate the likelihood functions of ***μ*** and ***ζ*** as multivariate normal distributions.

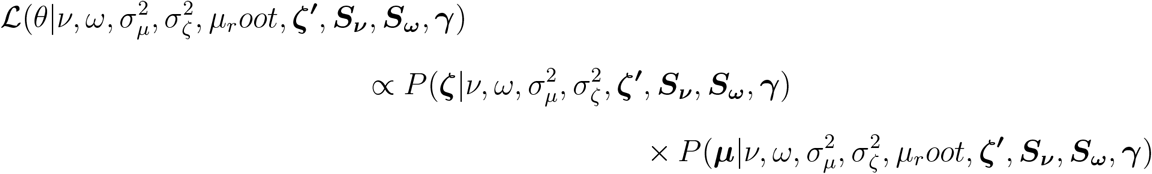

With ***μ, ζ, ζ*′** and ***γ*** being respectively observed means and log standard deviations, ancestral log standard deviations and branch lengths.

### Parameter estimation

The Bayesian estimation of the asymmetry (*ν*), the displacement (*ω*) and the evolutionary rates 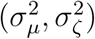 is dependant on the approximation of ancestral states (***ζ*′**, *μ*_root_) and switch parameters (***S***_*ν*_, ***S***_*ω*_). These parameters can be estimated using Gibbs sampling by calculating their conditional distributions on *ν, ω*, 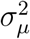 and 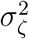. For our example highlighted in green (Fig. A1), in order to get the conditional distribution of *ζ*_5_ on *ζ*_1_, *ζ*_2_, *ν* and *ω*, we can add the expressions from (A3) and (A4) :

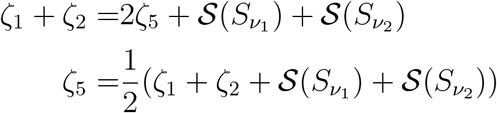

Because 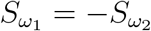, the sum 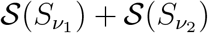 does not vary according to 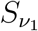. It is therefore a constant for a fixed *ν* and *ω* across all nodes that we note C = 𝒮(−1) + 𝒮(1)

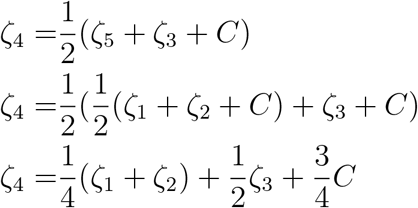

We can then write *ζ*_5_ and *ζ*_4_ as a linear combination of normal distributions:

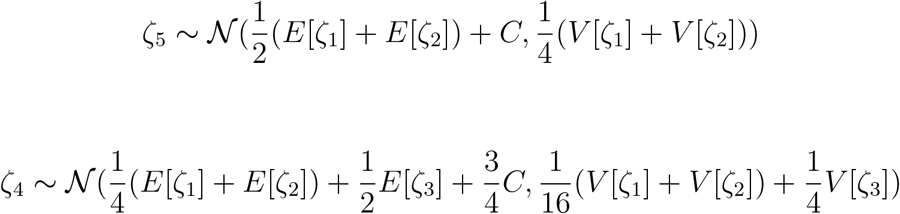

Using it we can show that for any node *k, ζ*_*k*_ is a linear combination of its descendants’ *ζ*. Therefore for every node *k* with *I*_*k*_ extant descendants and *J*_*k*_ descending edges, we have:

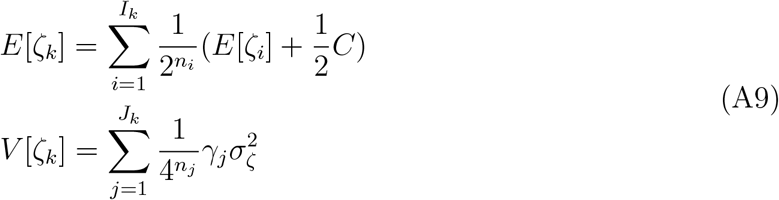

with *n*_*i*_ and *n*_*j*_ respectively the number of nodes between the node *k* and *i, j*. For every node with direct descendants *a* and *b* we can also write:

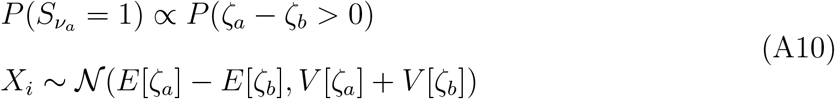

After the sampling of ***ζ*′** and ***S***_*ν*_ in their respective conditional distributions, we can calculate the conditional distribution of ***μ*′** and ***S***_*ω*_. In our example, we want to calculate the conditional distribution of *μ*_5_ on *μ*_1_, *μ*_2_, *ν, ω*, 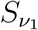 and 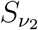. In order to get it we want to write an expression of μ_5_ as a weighted mean of *μ*_1_ and *μ*_2_. The weights represent the influence of the ancestral distribution inheritance on the descendants means:

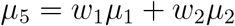

with *w*_1_ + *w*_2_ = 1. Using (A3) and (A4), we get:

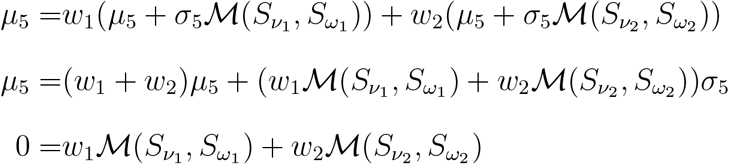

Then we get:

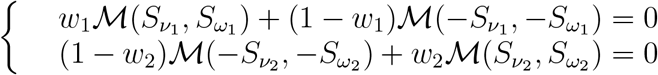

Which simplifies to :

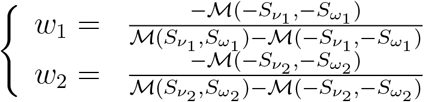

for *ν* > 0 and *ω* > 0. For *ω* = 0 or *ν* = 0 the descendants’ means or variances variances are equal so we have 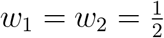. We then have:

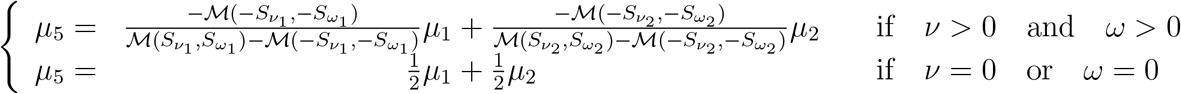

We can show that *w*_1_ and *w*_2_ are constant for a fixed *ν, ω* and 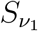, so we can calculate the conditional distribution of *μ*_5_ as a linear combination of *μ*_1_ and *μ*_2_ which are normally distributed:

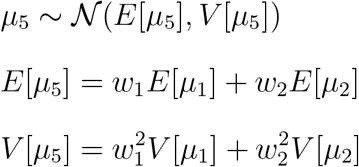

Similarly, for *μ*_4_ we find:

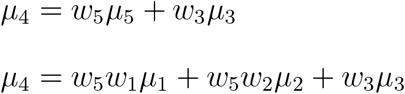

We can show that for any node *k, μ*_*k*_ is a linear combination of its descendants’ *μ*. Therefore for any node *k* with *I*_*k*_ descendants and *J*_*k*_ descending branches, we have:

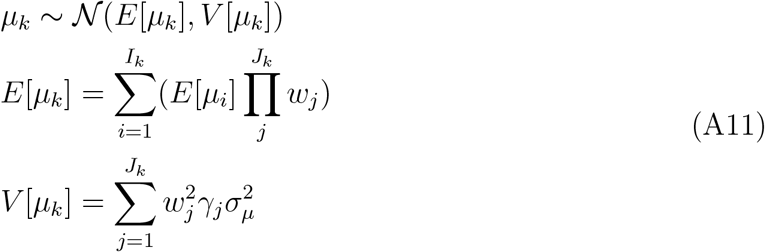

For every node with direct descendants a and b we can also write:

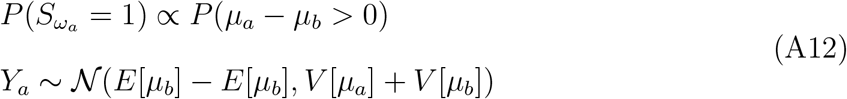

### Supplementary figures

**Fig. A2.**
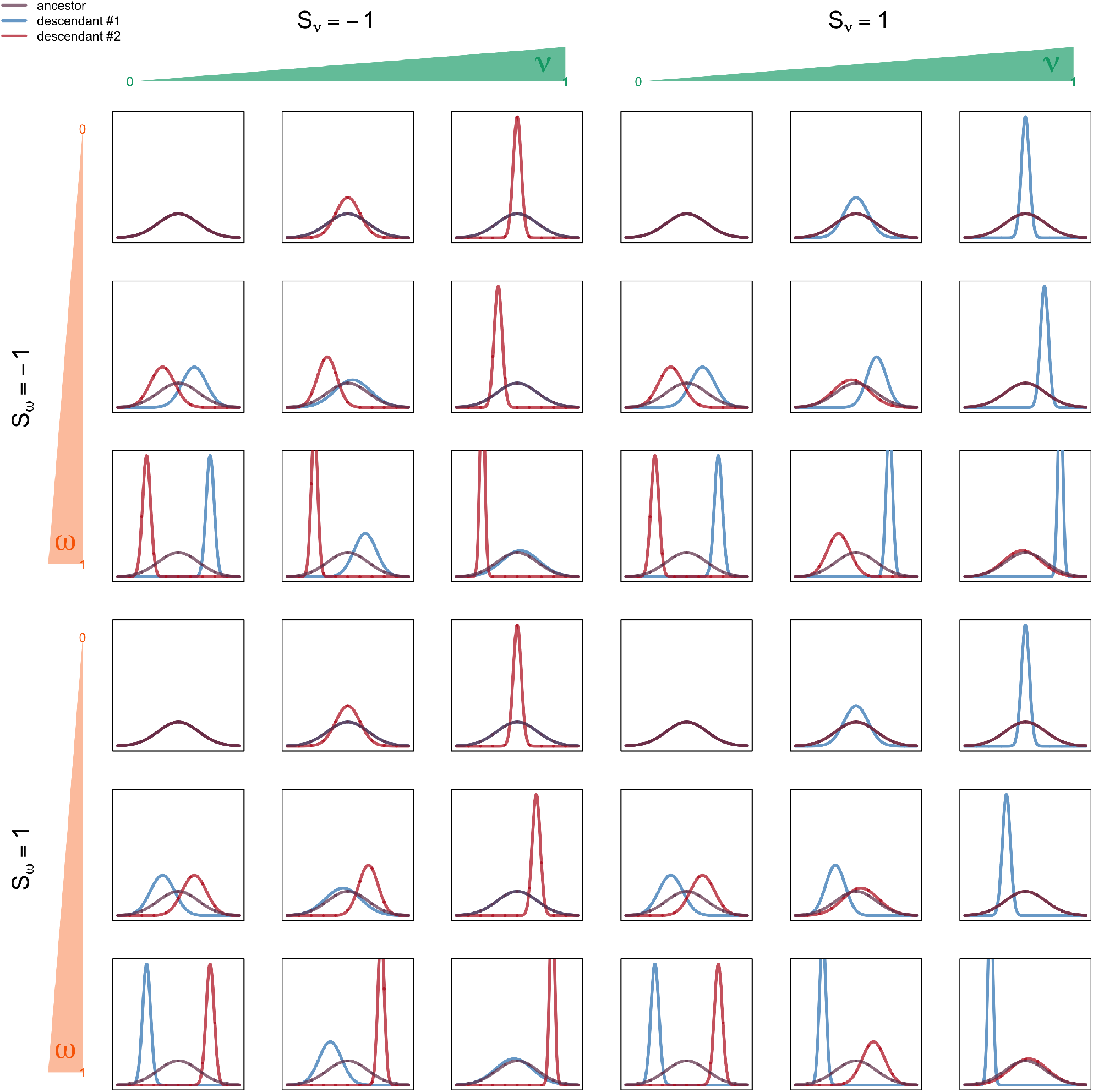
Alternative scenarios covered by the Asymmetric and Displaced Inheritance Process

**Fig. A3.**
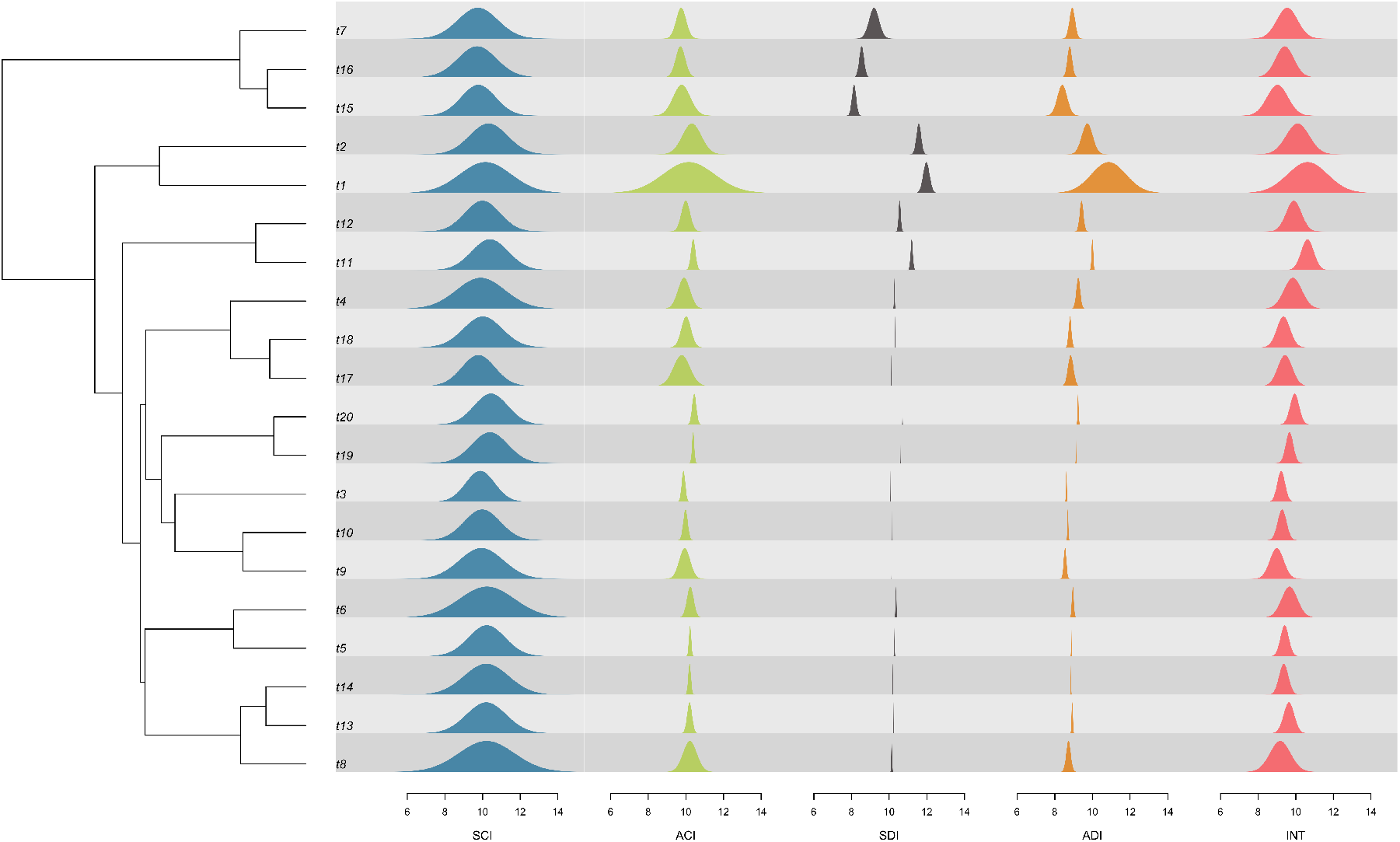
Example of simulated distributions with the different modes of trait distribution inheritance at the time of speciation and 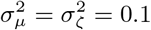. SCI: Symmetric and conserved inheritance *ν* = 0, *ω* = 0, ACI: Asymmetric and Conserved inheritance *ν* = 0.5, *ω* = 0, SDI: Symmetric and Displaced inheritance *ν* = 0, *ω* = 0.5, ADI: Asymmetric and Displaced inheritance *ν* = 0.5, *ω* = 0.5, INT: intermediate scenario *ν* = 0.2, *ω* = 0.2

**Fig. A4.**
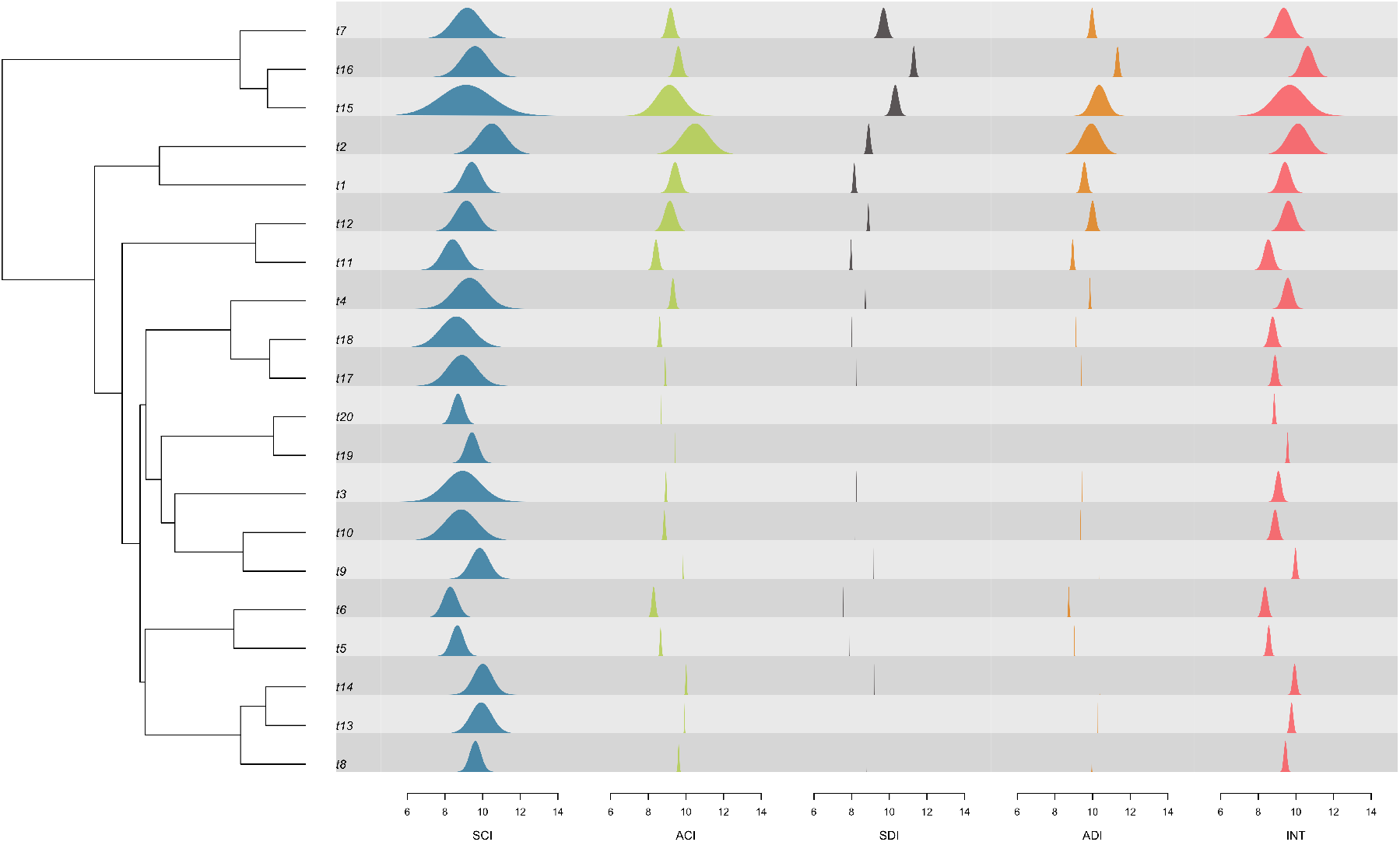
Example of simulated distributions with the different modes of trait distribution inheritance at the time of speciation and 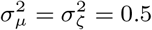. SCI: Symmetric and conserved inheritance *ν* = 0, *ω* = 0, ACI: Asymmetric and Conserved inheritance *ν* = 0.5, *ω* = 0, SDI: Symmetric and Displaced inheritance *ν* = 0, *ω* = 0.5, ADI: Asymmetric and Displaced inheritance *ν* = 0.5, *ω* = 0.5, INT: intermediate scenario *ν* = 0.2, *ω* = 0.2

**Fig. A5.**
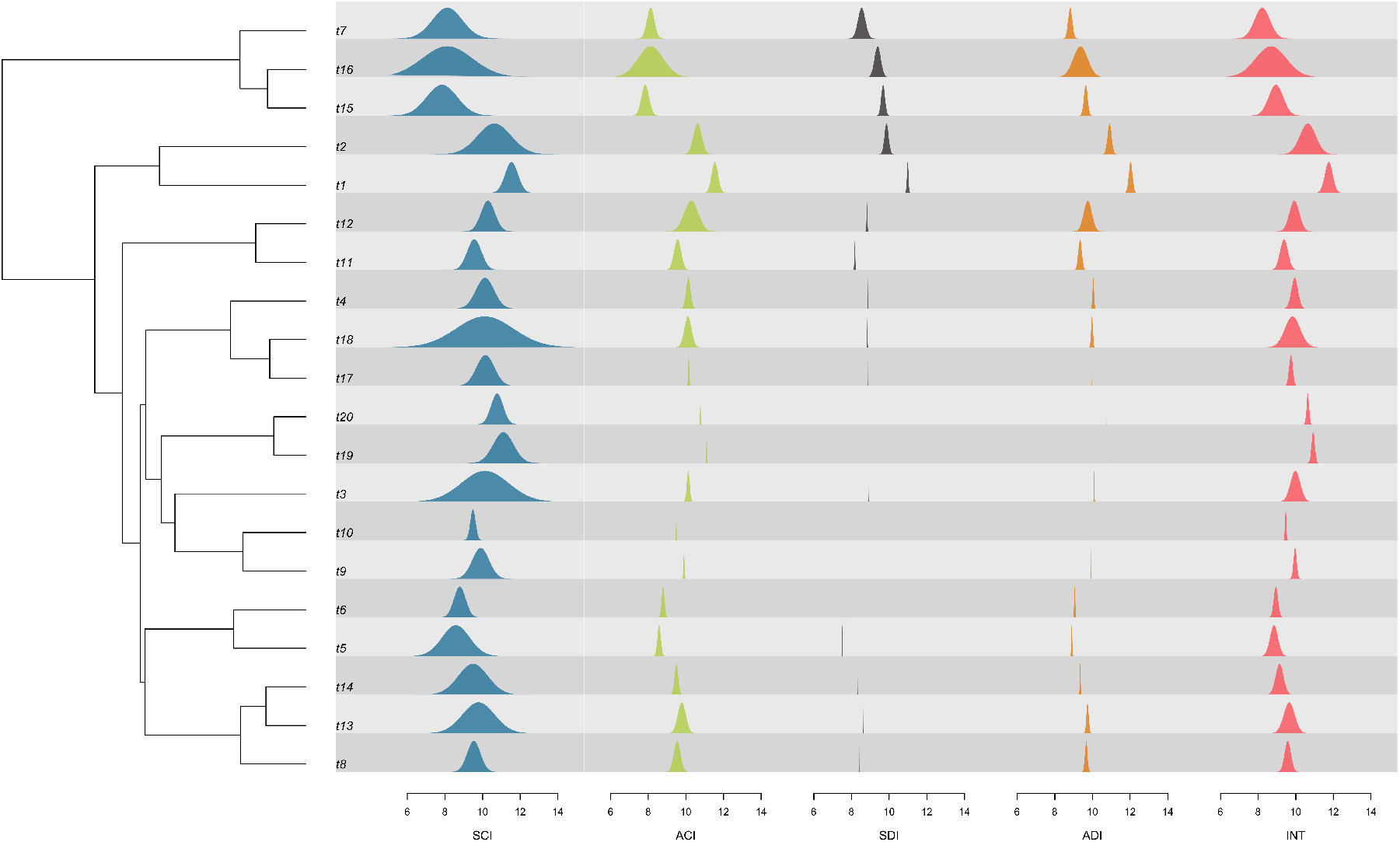
Example of simulated distributions with the different modes of trait distribution inheritance at the time of speciation and 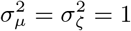. SCI: Symmetric and conserved inheritance *ν* = 0, *ω* = 0, ACI: Asymmetric and Conserved inheritance *ν* = 0.5, *ω* = 0, SDI: Symmetric and Displaced inheritance *ν* = 0, *ω* = 0.5, ADI: Asymmetric and Displaced inheritance *ν* = 0.5, *ω* = 0.5, INT: intermediate scenario *ν* = 0.2, *ω* = 0.2

**Fig. A6.**
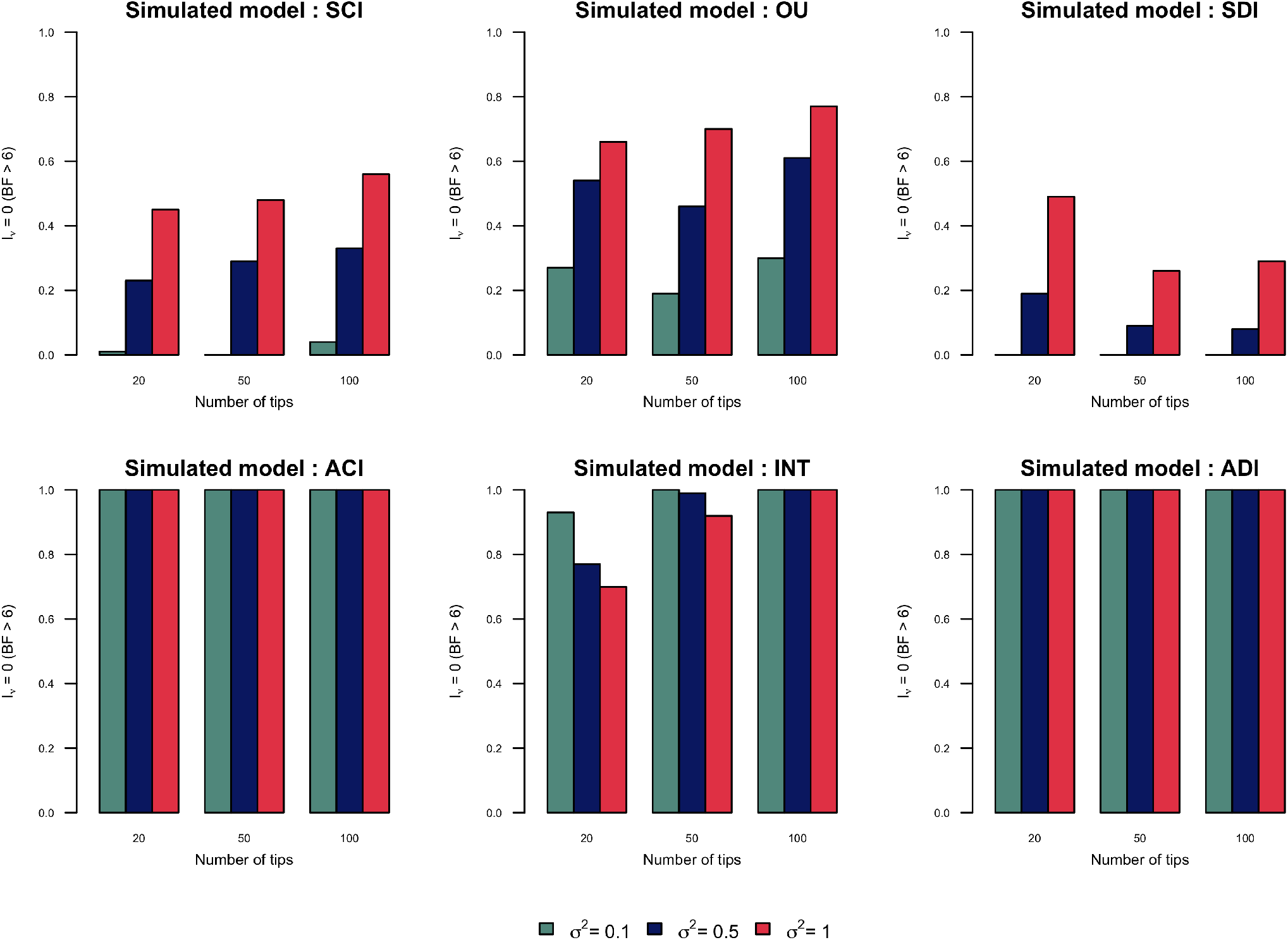
Proportion of simulations for which asymmetry has been rejected in function of the number of tips and evolutionary rates. SCI: Symmetric and conserved inheritance *ν* = 0, *ω* = 0, ACI: Asymmetric and Conserved inheritance *ν* = 0.5, *ω* = 0, SDI: Symmetric and Displaced inheritance *ν* = 0, *ω* = 0.5, ADI: Asymmetric and Displaced inheritance *ν* = 0.5, *ω* = 0.5, INT: intermediate scenario *ν* = 0.2, *ω* = 0.2

**Fig. A7.**
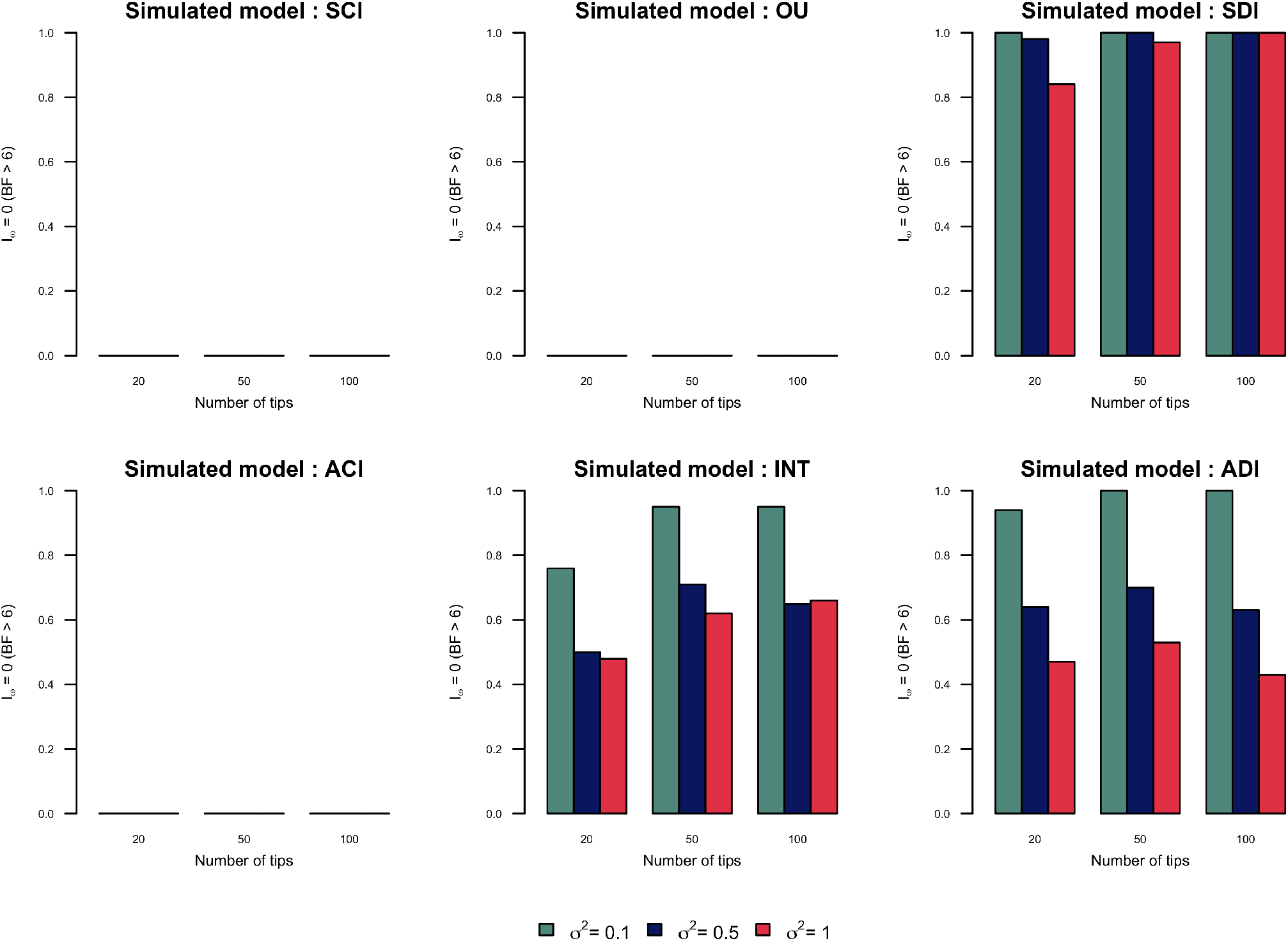
Proportion of simulations for which displacement has been rejected in function of the number of tips and evolutionary rates. SCI: Symmetric and conserved inheritance *ν* = 0, *ω* = 0, ACI: Asymmetric and Conserved inheritance *ν* = 0.5, *ω* = 0, SDI: Symmetric and Displaced inheritance *ν* = 0, *ω* = 0.5, ADI: Asymmetric and Displaced inheritance *ν* = 0.5, *ω* = 0.5, INT: intermediate scenario *ν* = 0.2, *ω* = 0.2

**Fig. A8.**
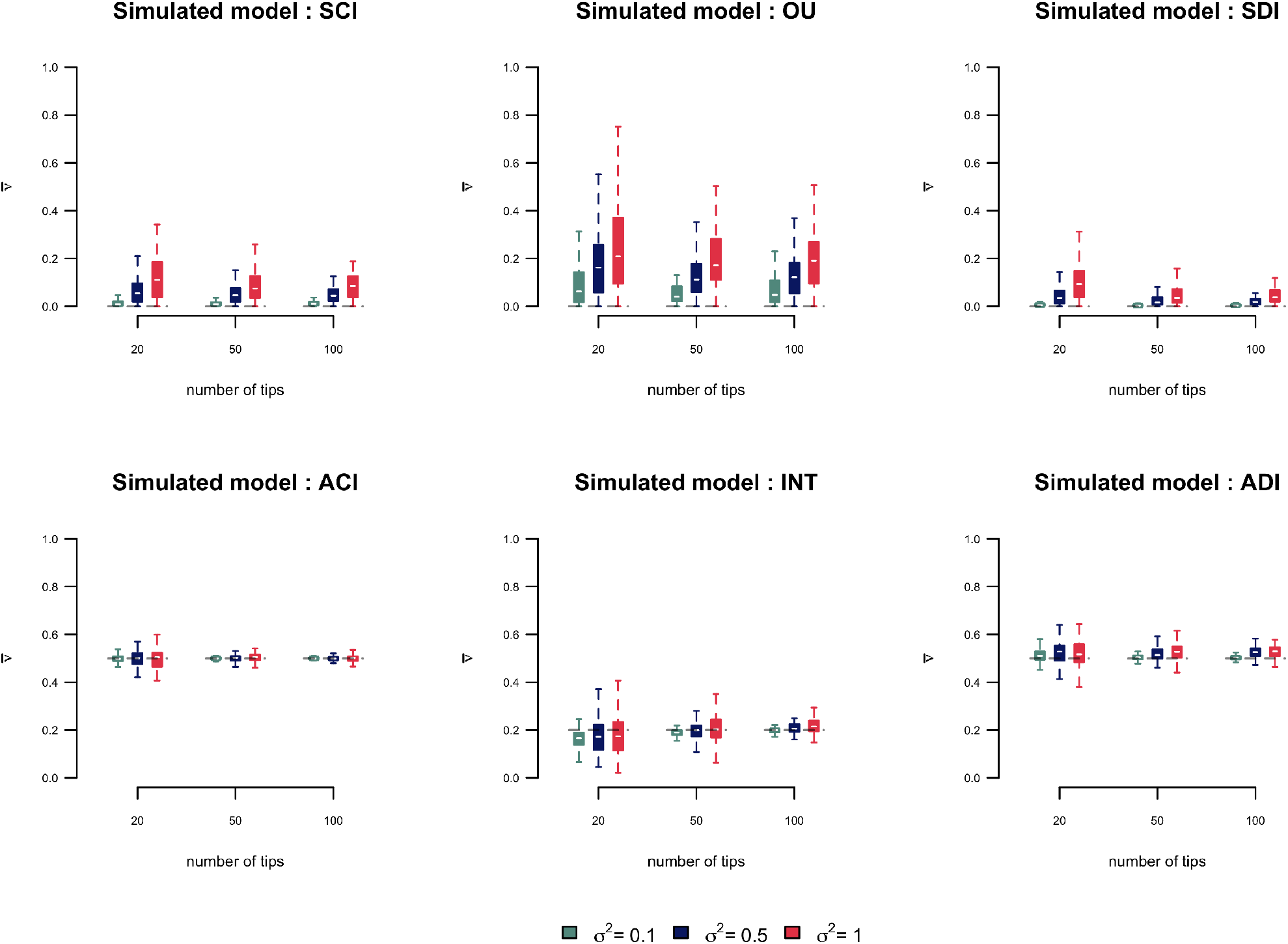
Distribution of mean estimates of *ν* from homogeneous simulations in function of the number of tips and evolutionary rates. SCI: Symmetric and conserved inheritance *ν* = 0, *ω* = 0, ACI: Asymmetric and Conserved inheritance *ν* = 0.5, *ω* = 0, SDI: Symmetric and Displaced inheritance *ν* = 0, *ω* = 0.5, ADI: Asymmetric and Displaced inheritance *ν* = 0.5, *ω* = 0.5, INT: intermediate scenario *ν* = 0.2, *ω* = 0.2

**Fig. A9.**
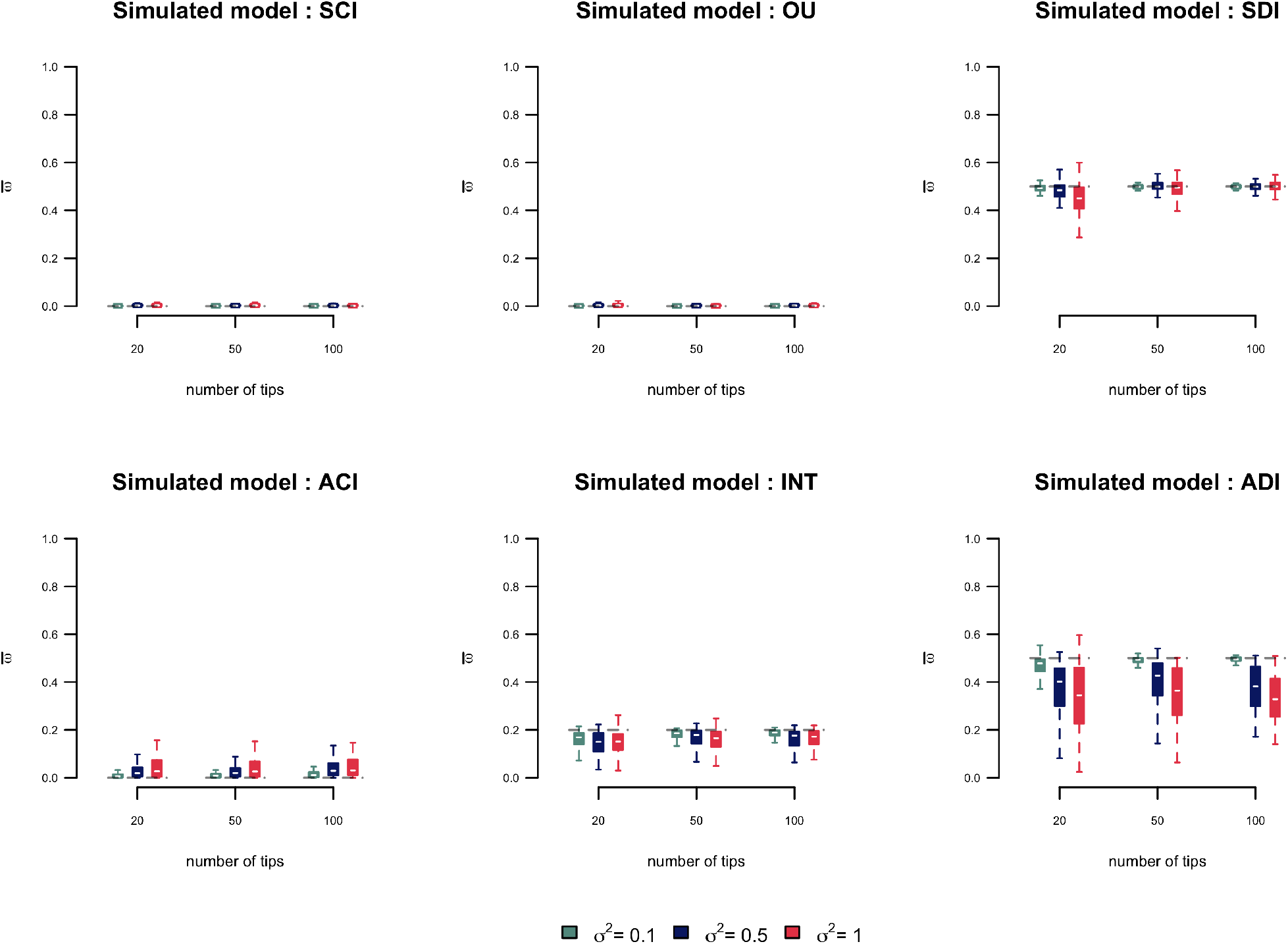
Distribution of mean estimates of *ω* from homogeneous simulations in function of the number of tips and evolutionary rates. SCI: Symmetric and conserved inheritance *ν* = 0, *ω* = 0, ACI: Asymmetric and Conserved inheritance *ν* = 0.5, *ω* = 0, SDI: Symmetric and Displaced inheritance *ν* = 0, *ω* = 0.5, ADI: Asymmetric and Displaced inheritance *ν* = 0.5, *ω* = 0.5, INT: intermediate scenario *ν* = 0.2, *ω* = 0.2

**Fig. A10.**
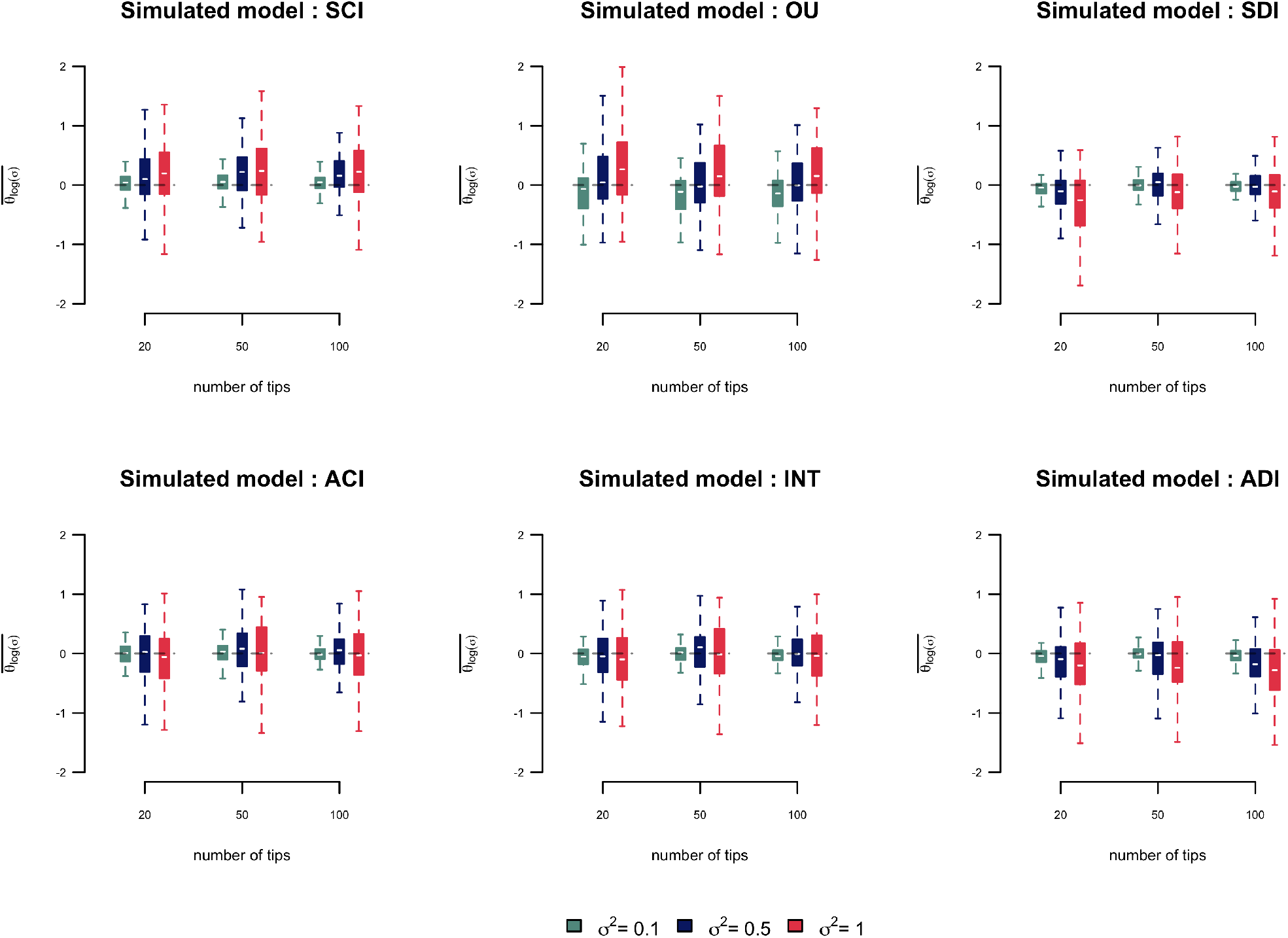
Distribution of mean estimates of *θ*_*ζ*_ from homogeneous simulations in function of the number of tips and evolutionary rates. SCI: Symmetric and conserved inheritance *ν* = 0, *ω* = 0, ACI: Asymmetric and Conserved inheritance *ν* = 0.5, *ω* = 0, SDI: Symmetric and Displaced inheritance *ν* = 0, *ω* = 0.5, ADI: Asymmetric and Displaced inheritance *ν* = 0.5, *ω* = 0.5, INT: intermediate scenario *ν* = 0.2, *ω* = 0.2

**Fig. A11.**
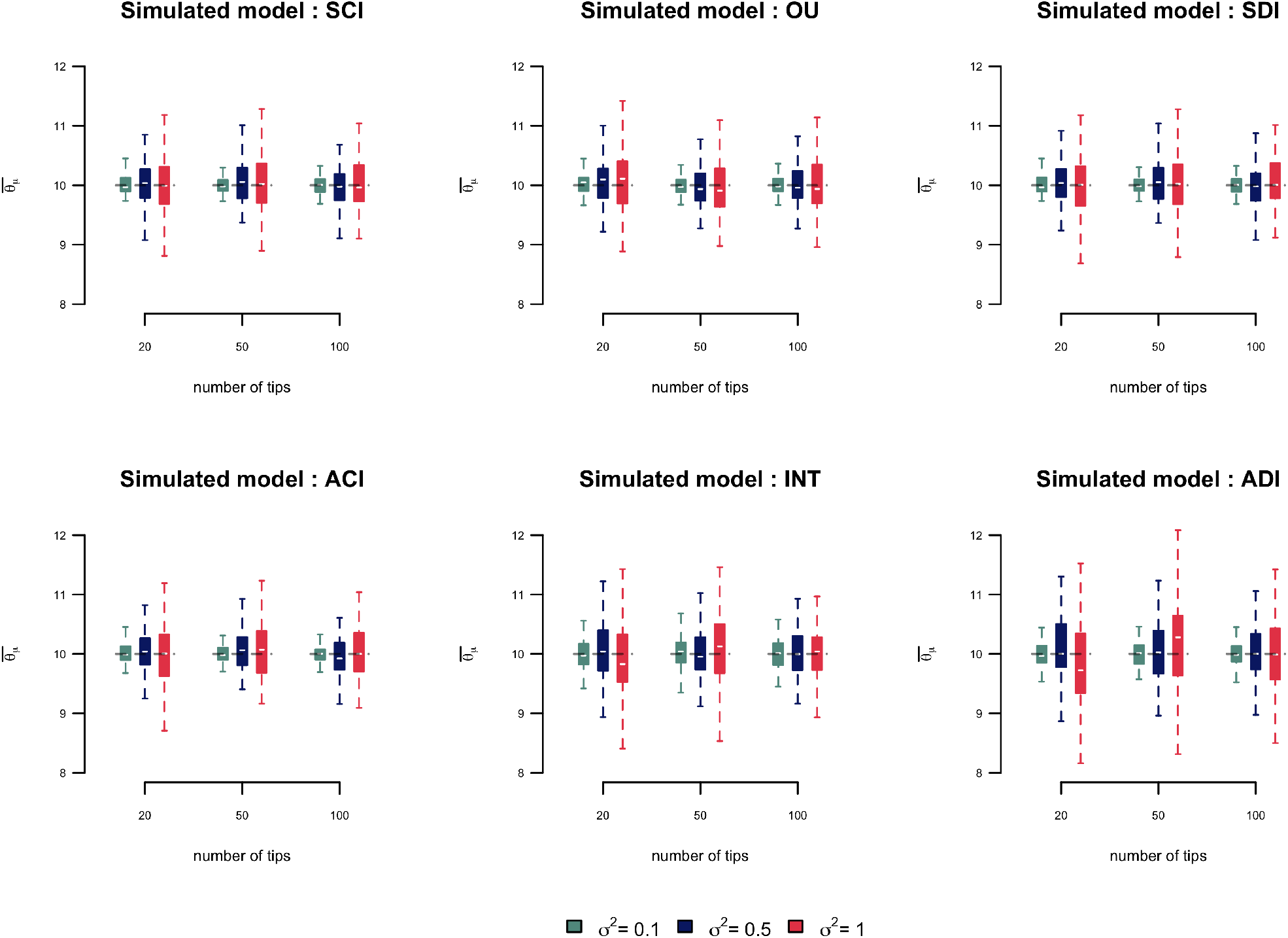
Distribution of mean estimates of *θ*_*μ*_ from homogeneous simulations in function of the number of tips and evolutionary rates. SCI: Symmetric and conserved inheritance *ν* = 0, *ω* = 0, ACI: Asymmetric and Conserved inheritance *ν* = 0.5, *ω* = 0, SDI: Symmetric and Displaced inheritance *ν* = 0, *ω* = 0.5, ADI: Asymmetric and Displaced inheritance *ν* = 0.5, *ω* = 0.5, INT: intermediate scenario *ν* = 0.2, *ω* = 0.2

**Fig. A12.**
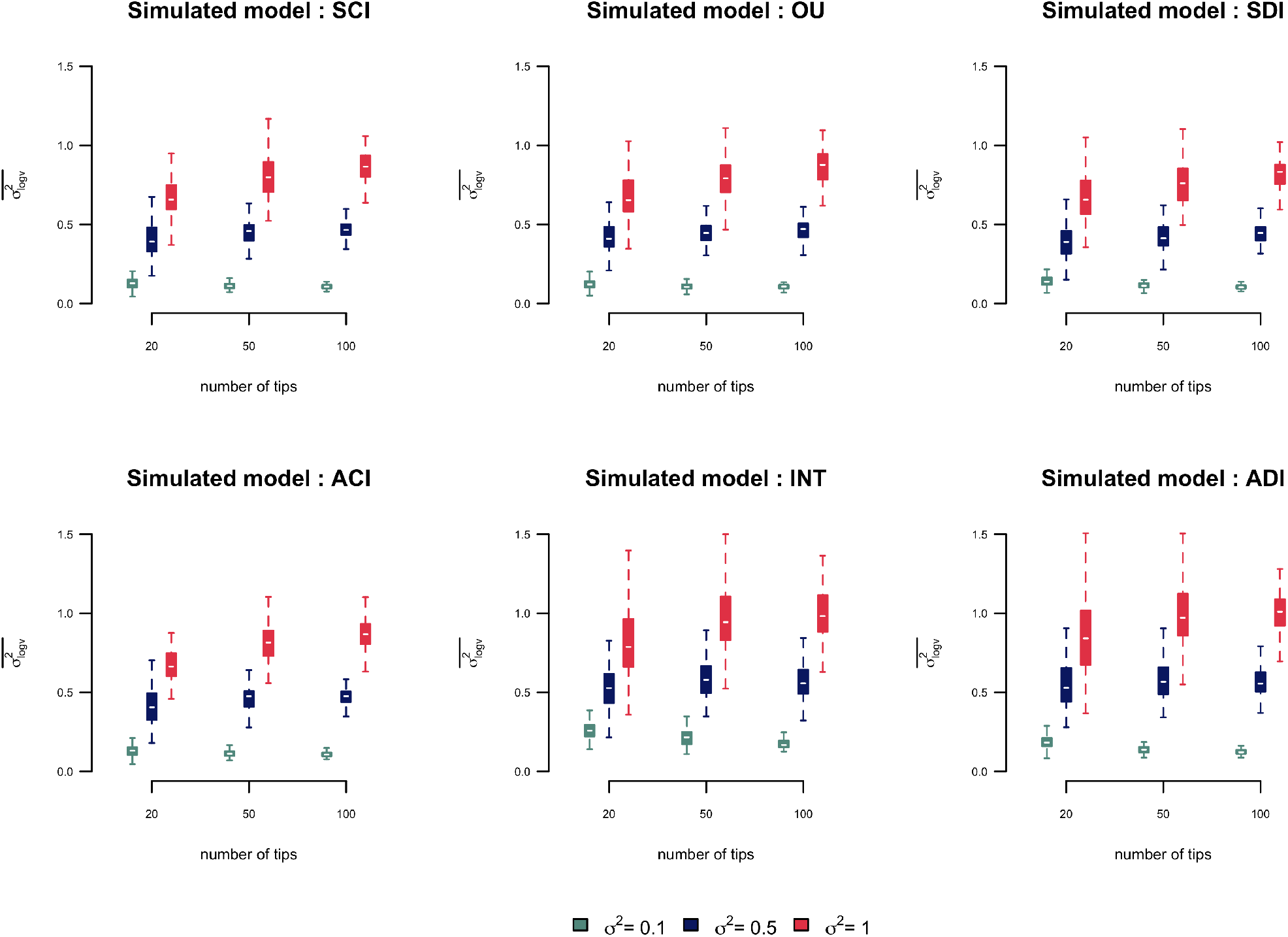
Distribution of mean estimates of 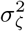 from homogeneous simulations in function of the number of tips and evolutionary rates. SCI: Symmetric and conserved inheritance *ν* = 0, *ω* = 0, ACI: Asymmetric and Conserved inheritance *ν* = 0.5, *ω* = 0, SDI: Symmetric and Displaced inheritance *ν* = 0, *ω* = 0.5, ADI: Asymmetric and Displaced inheritance *ν* = 0.5, *ω* = 0.5, INT: intermediate scenario *ν* = 0.2, *ω* = 0.2

**Fig. A13.**
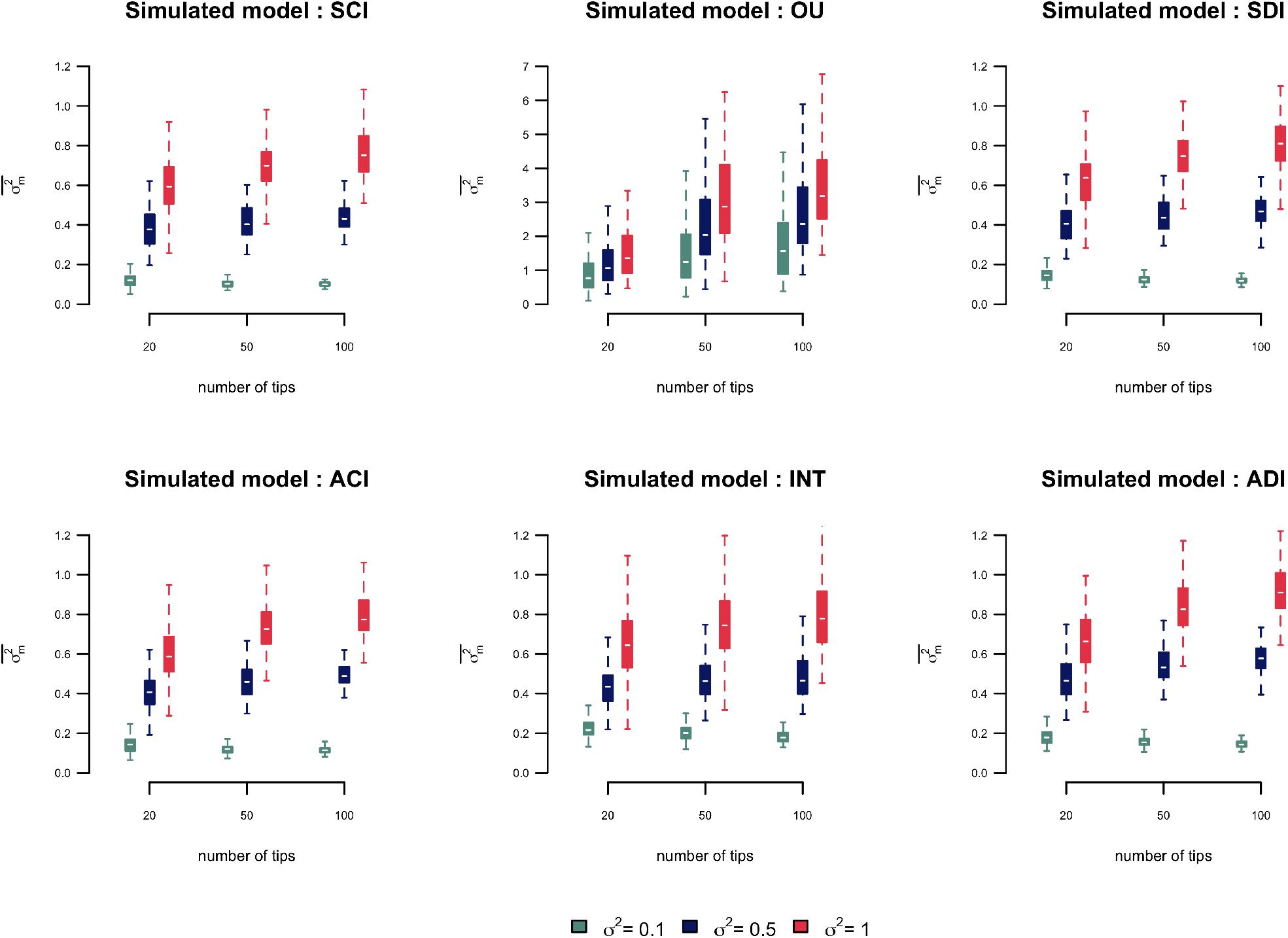
Distribution of mean estimates of 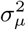 from homogeneous simulations in function of the number of tips and evolutionary rates. SCI: Symmetric and conserved inheritance *ν* = 0, *ω* = 0, ACI: Asymmetric and Conserved inheritance *ν* = 0.5, *ω* = 0, SDI: Symmetric and Displaced inheritance *ν* = 0, *ω* = 0.5, ADI: Asymmetric and Displaced inheritance *ν* = 0.5, *ω* = 0.5, INT: intermediate scenario *ν* = 0.2, *ω* = 0.2

**Fig. A14.**
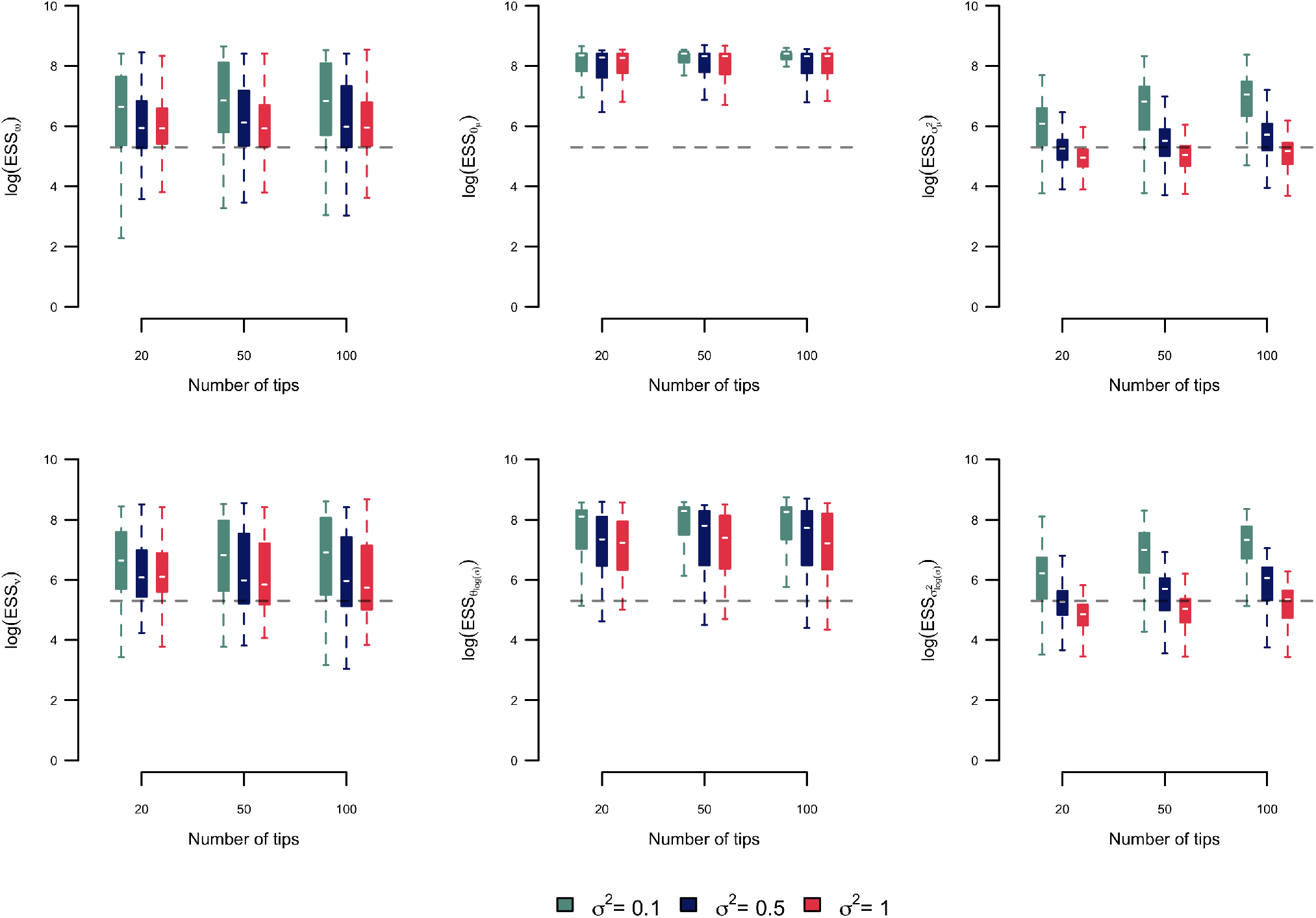
log Estimated sample size (ESS) of ABM parameters estimated from the outputs of the MCMC algorithm applied on the first dataset. The dashed lines represent an ESS = 200

**Fig. A15.**
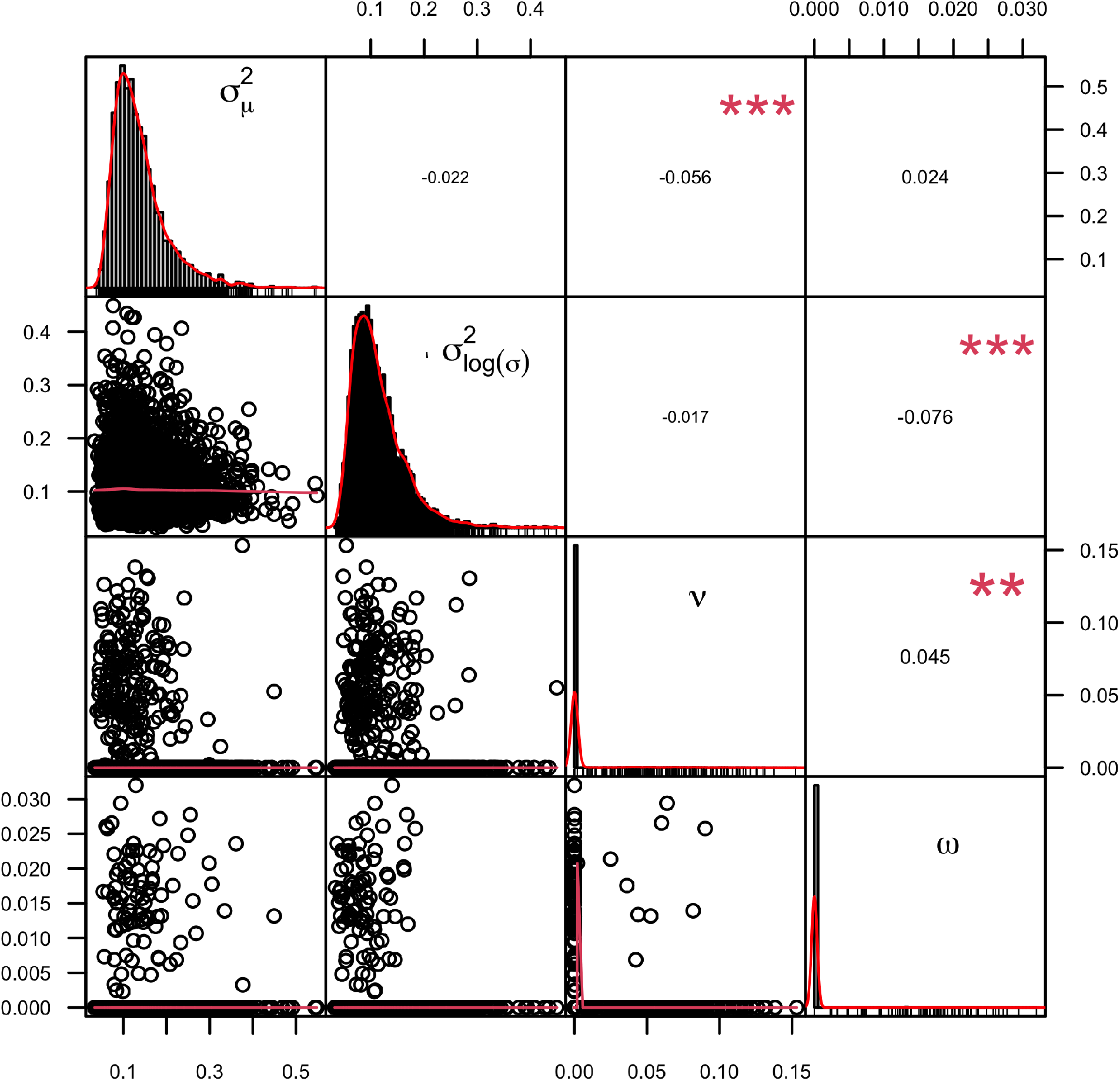
Correlation between parameters in one example simulation with *n* = 20 *ν* = 0, *ω* = 0 and 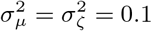 Right panels represent the statistic and p-value of Spearman ranked test. Diagonals represent the posterior distributions

**Fig. A16.**
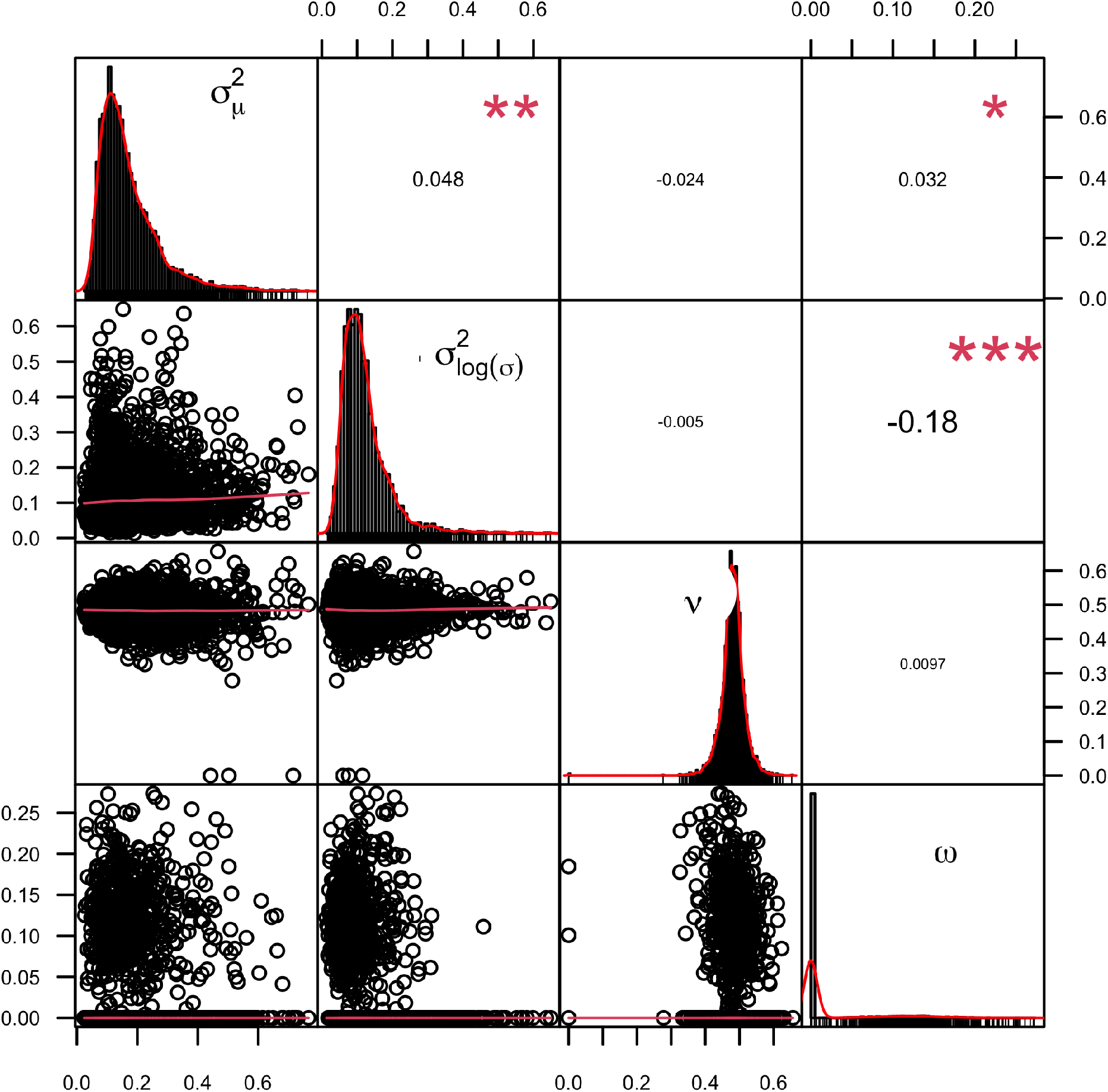
Correlation between parameters in one example simulation with *n* = 20 *ν* = 0.5, *ω* = 0 and 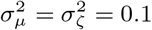 Right panels represent the statistic and p-value of Spearman ranked test. Diagonals represent the posterior distributions

**Fig. A17.**
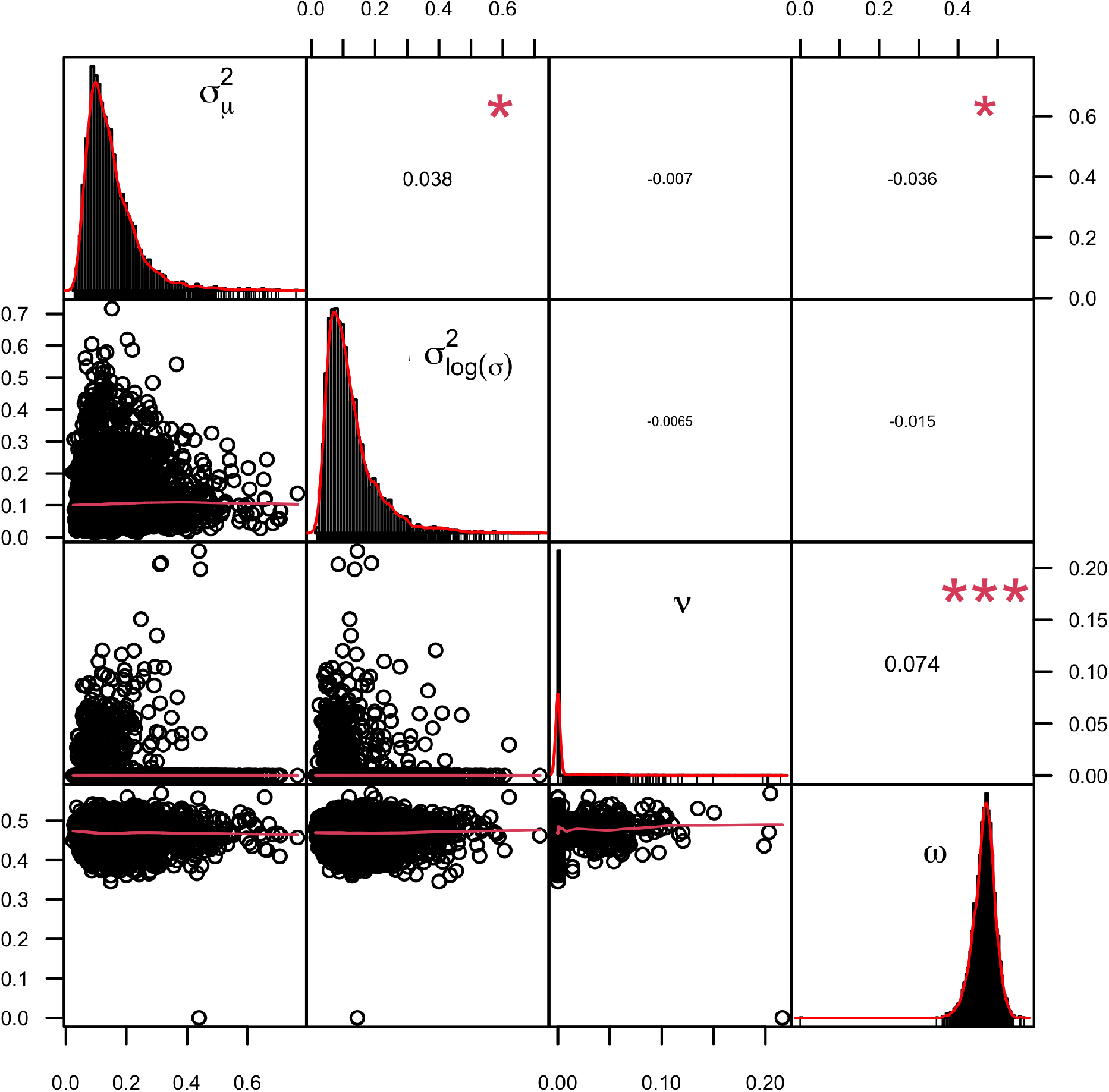
Correlation between parameters in one example simulation with *n* = 20 *ν* = 0, *ω* = 0.5 and 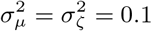 Right panels represent the statistic and p-value of Spearman ranked test. Diagonals represent the posterior distributions

**Fig. A18.**
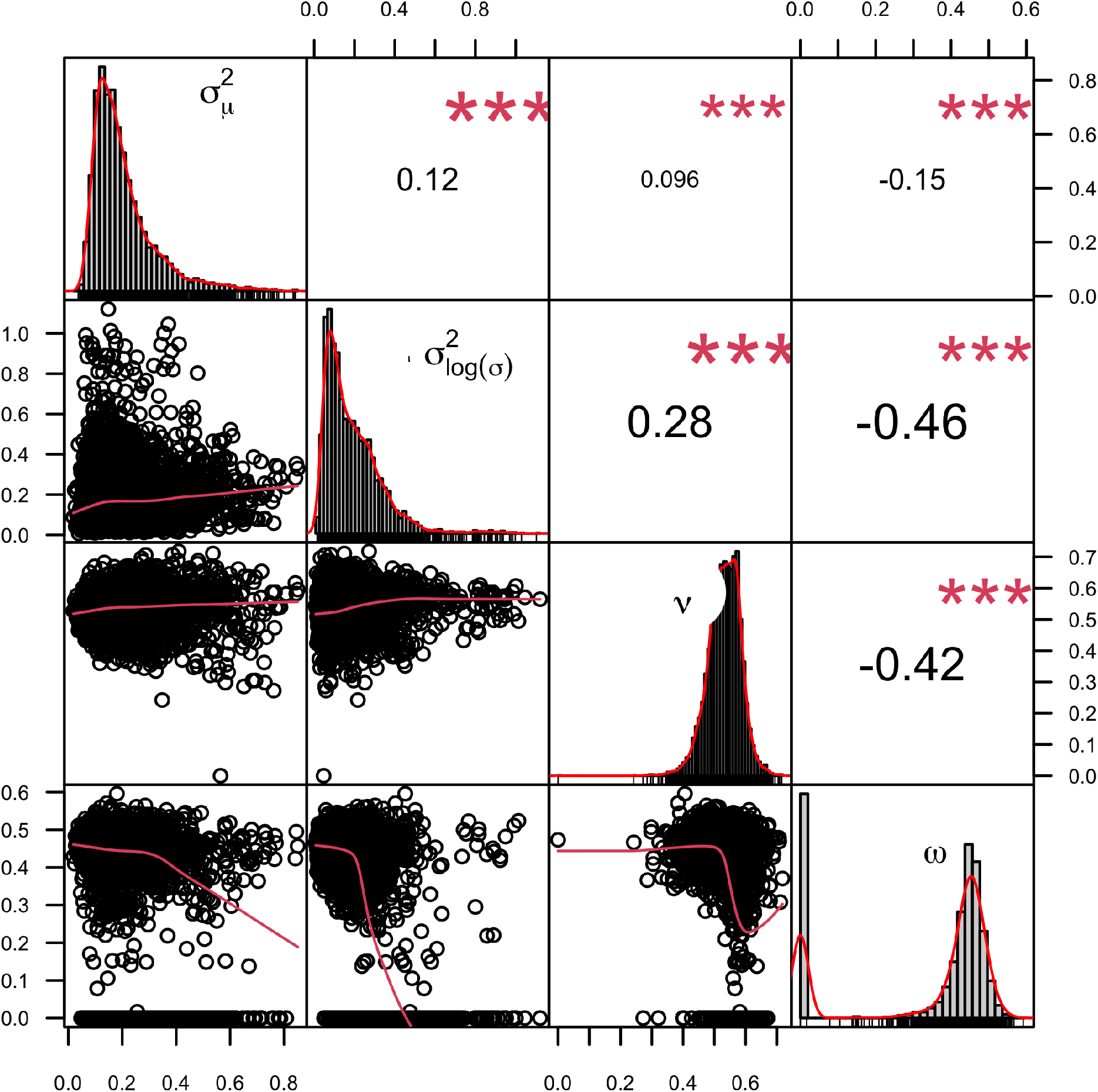
Correlation between parameters in one example simulation with *n* = 20 *ν* = 0.5, *ω* = 0.5 and 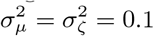 Right panels represent the statistic and p-value of Spearman ranked test. Diagonals represent the posterior distributions

**Fig. A19.**
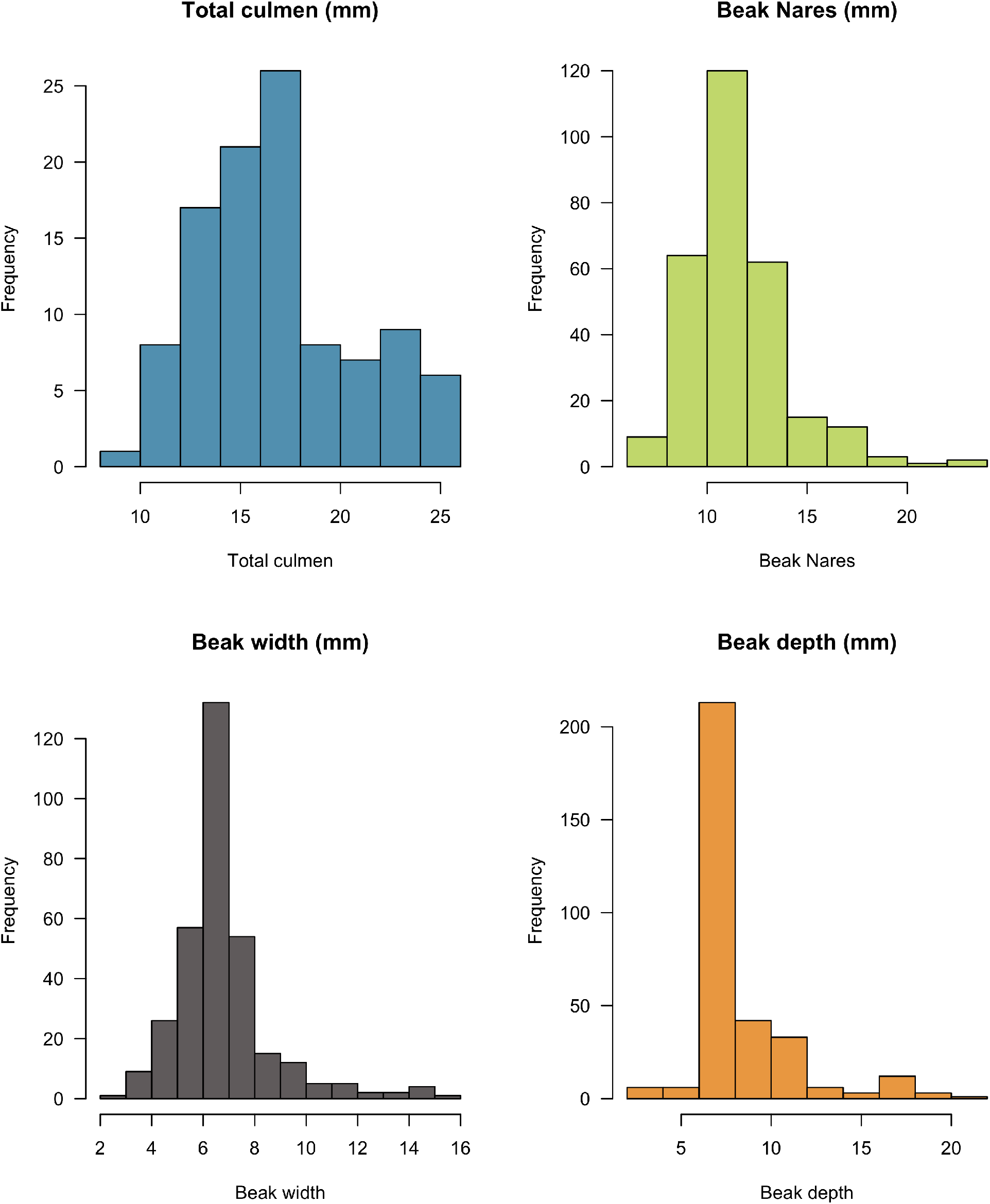
Distribution of individual phenotypic measurements in Coerebinae

**Fig. A20.**
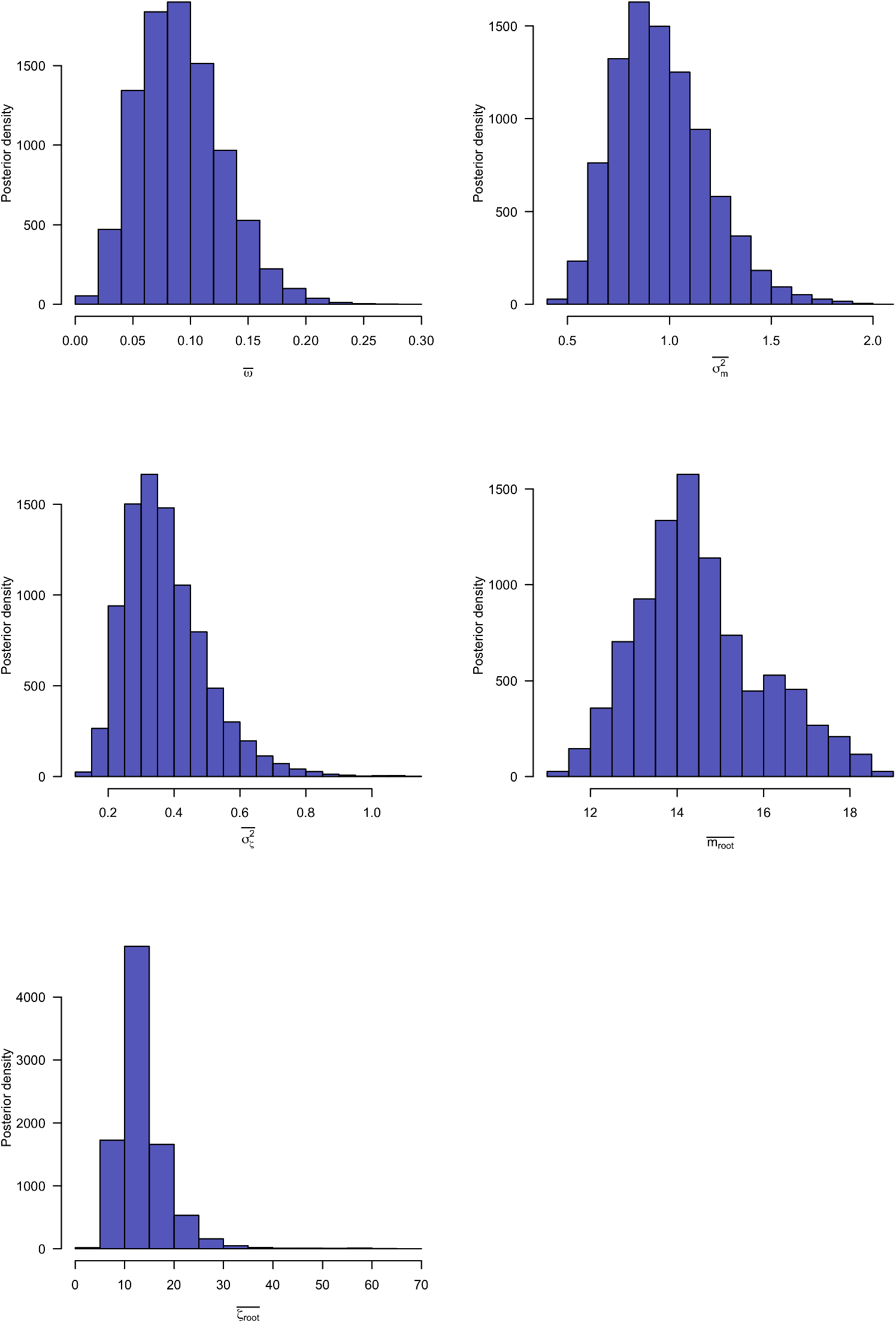
Posterior distribution of estimated parameters using the ABM for total culmen

**Fig. A21.**
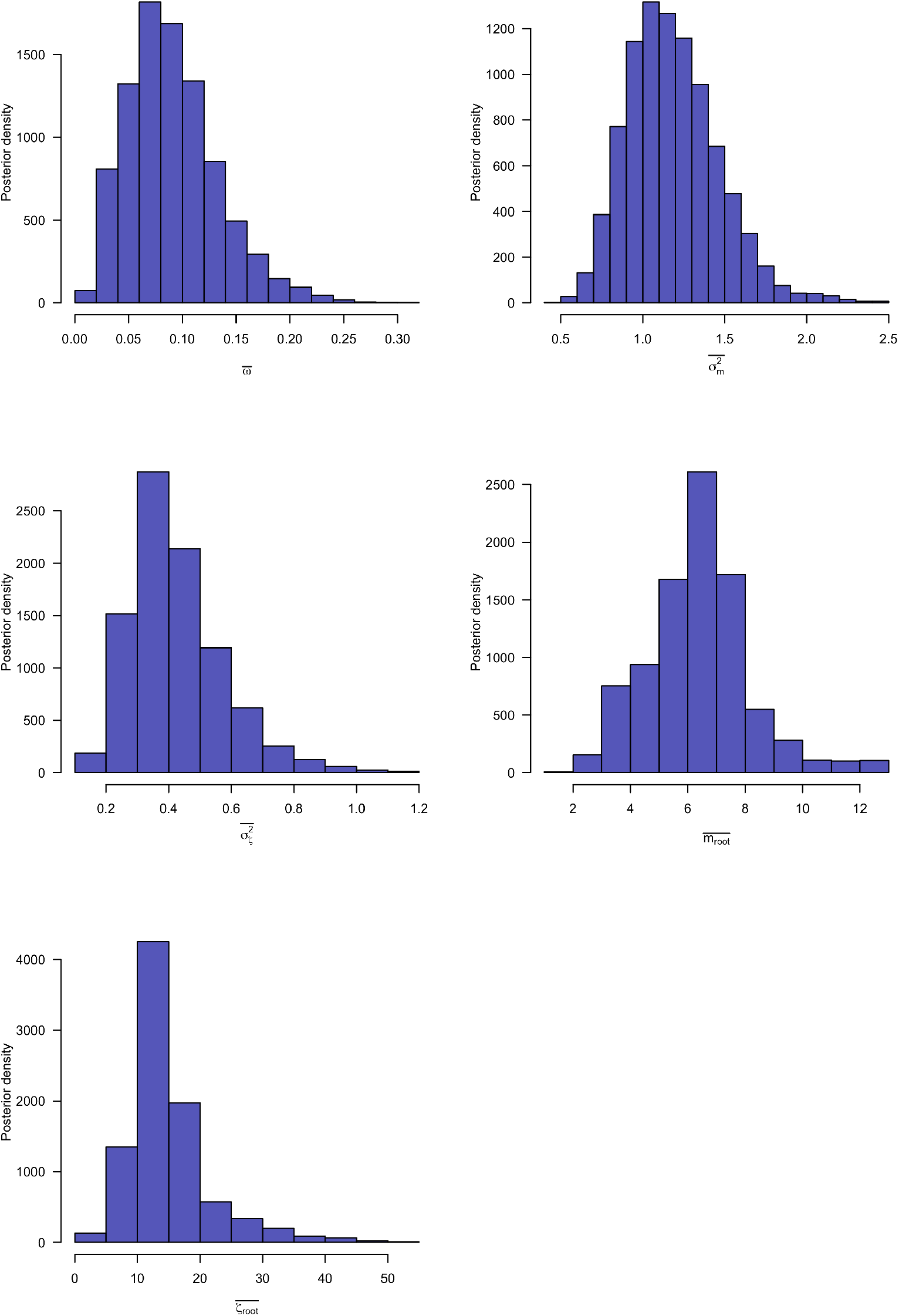
Posterior distribution of estimated parameters using the ABM for bill depth

**Fig. A22.**
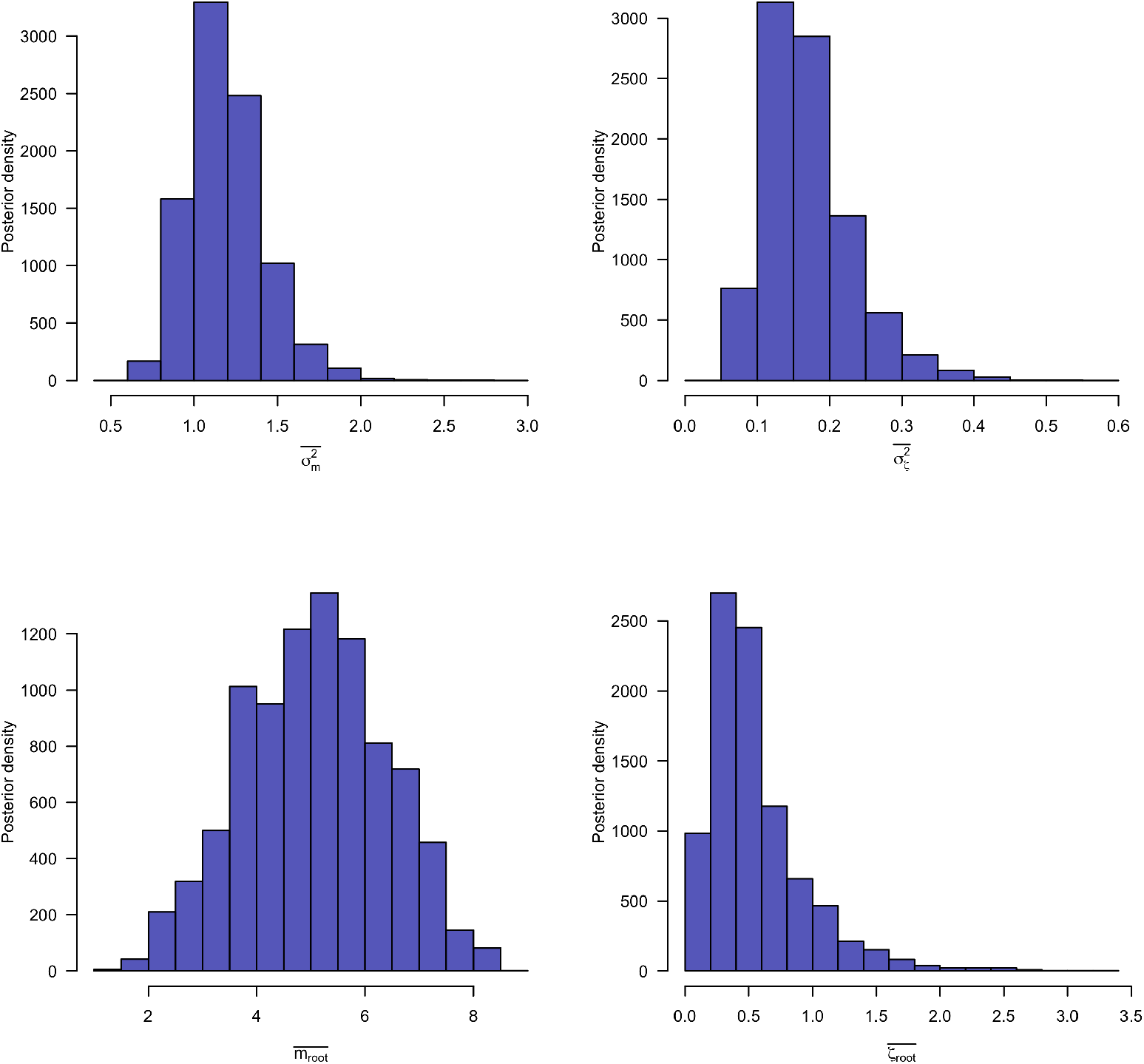
Posterior distribution of estimated parameters using the ABM for bill width

**Fig. A23.**
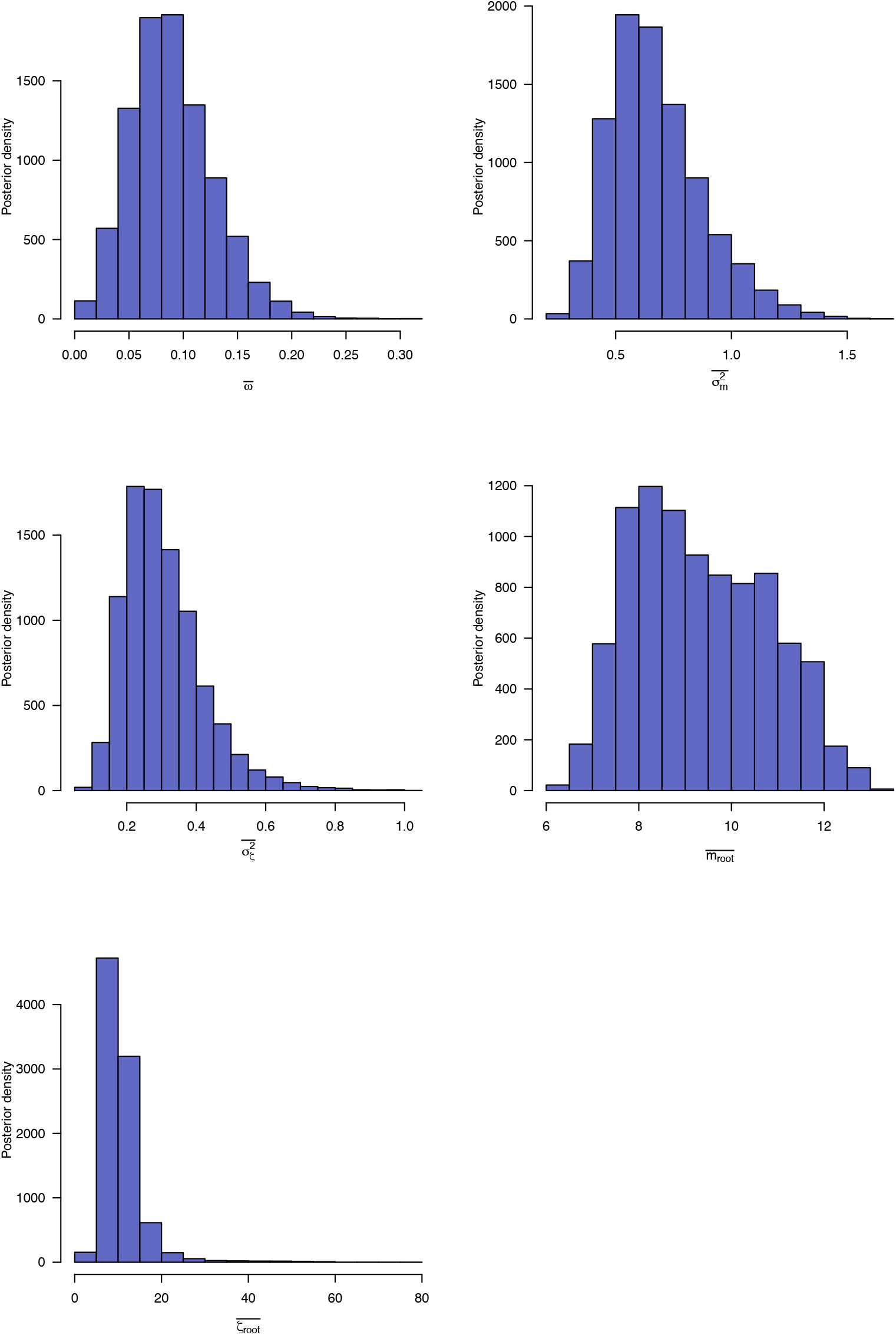
Posterior distribution of estimated parameters using the ABM for bill nares

## Notes

https://doi.org/10.5061/dryad.q573n5tns

### Competing Interest Statement

The authors have declared no competing interest.

